# Confronting pitfalls of AI-augmented molecular dynamics using statistical physics

**DOI:** 10.1101/2020.06.11.146985

**Authors:** Shashank Pant, Zachary Smith, Yihang Wang, Emad Tajkhorshid, Pratyush Tiwary

## Abstract

Artificial intelligence (AI)-based approaches have had indubitable impact across the sciences through the ability to extract relevant information from raw data. Recently AI has also seen use for enhancing the efficiency of molecular simulations, wherein AI derived slow modes are used to accelerate the simulation in targeted ways. However, while typical fields where AI is used are characterized by a plethora of data, molecular simulations per-construction suffer from limited sampling and thus limited data. As such the use of AI in molecular simulations can suffer from a dangerous situation where the AI-optimization could get stuck in spurious regimes, leading to incorrect characterization of the reaction coordinate (RC) for the problem at hand. When such an incorrect RC is then used to perform additional simulations, one could start to deviate progressively from the ground truth. To deal with this problem of spurious AI-solutions, here we report a novel and automated algorithm using ideas from statistical mechanics. It is based on the notion that a more reliable AI-solution will be one that maximizes the time-scale separation between slow and fast processes. To learn this time-scale separation even from limited data, we use a maximum caliber-based framework. We show the applicability of this automatic protocol for 3 classic benchmark problems, namely the conformational dynamics of a model peptide, ligand-unbinding from a protein, and folding/unfolding energy landscape of the C-terminal domain of protein G. We believe our work will lead to increased and robust use of trustworthy AI in molecular simulations of complex systems.

## Introduction

With the development of more accurate force fields and powerful computers, molecular dynamics (MD) has become a ubiquitous tool to study complex structural, thermodynamic and kinetic processes of real-world systems across disciplines. However, the predictive capacity of the methodology is limited by the large time-scale gap between the conformational dynamics of the complex processes of interest and the short periods accessible to it.^1,2^ This disparity is mostly attributed to the rough energy landscape typically characterized by numerous energy minima with hard to cross barriers between them,^1,3,4^ which trap the system in metastable states, leading to an incomplete sampling of the configuration space. Comprehensive sampling of the configuration space not only provides high temporal and spatial resolutions of the complex process but also allows us to compute converged thermodynamic properties, sample physiologically relevant molecular conformations, and explore complex motions critical to biological and chemical processes such as protein folding, ligand binding, energy transfer, and countless others.^5–13^

To overcome the limitations of time-scales and accurately characterize such complex landscapes, a plethora of enhanced sampling techniques have been developed. We can broadly divide these methods into: (1) tempering based, and (2) collective variable (CV) or reaction coordinate (RC) based,^4^ either of which can then also be coupled with multiple replica based exchange schemes. In tempering based methods, the underlying landscape is sampled by either modifying the temperature and/or Hamiltonian of the system through approaches like temperature replica exchange, simulated annealing, and accelerated MD.^14–21^ On the other hand, CV based methods involve enhancing fluctuations along pre-defined low-dimensional modes, through biased sampling approaches like metadynamics,^22–24^ umbrella-sampling (US),^25^ adaptive biasing force (ABF)^26–30^ and many others.^27,31–33^ Although CV-based methods can be computationally more efficient than tempering-based approaches, given a poor choice of low-dimensional modes (a non-trivial task to intuit for complex systems), CV biasing can fail miserably.^34^ Indeed, one could also argue that one way to make tempering methods more efficient, is to select a specific part of the system, akin to a CV, which is then subjected to the tempering protocol.^35,36^

Artificial intelligence (AI) potentially provides a systematic means to differentiate signal from noise in generic data, and thus discover relevant CVs to accelerate the simulations.^37–41^ A number of such AI-based approaches have been proposed recently^37–39,42,43^ and remain the subject of extensive research. A common underlying theme in these methods is to exploit AI tools to gradually uncover the underlying effective geometry, parametrize it on-the-fly, and exploit it to bias the design of experiments with the MD simulator by emphasizing informative configuration space areas that have not been explored before. This iterative MD-AI procedure is repeated until desired sampling has been achieved. Conceptually, these approaches effectively restrain the 3*N*-dimensional space to a very small number of dimensions (typically 1 or 2) which encode all the relevant slow dynamics in the system, effectively discarding the remaining fast dynamics. Every round of AI estimates the slow modes given sampling so far, and this information is used to launch new biased rounds of simulations. Biasing along the slow modes leads to increased exploration, which can then be used in another round of AI to estimate the relevant slow modes even more accurately. The use of standard reweighting procedures can then recover unbiased thermodynamic and kinetic information from the AI-augmented MD trajectories so obtained.

However, there is a fundamental problem in such an approach. Most AI tools are designed for data-rich systems. It has been argued^44–47^ that given good quality training data and with a neural network with infinitely many parameters, the objective function for associated stochastic gradient optimization schemes is convex. However in enhanced MD, we are per construction in a poorly sampled, data-sparse regime, and moreover, it is impractical to use a dense network with too many parameters. The AI optimization function is therefore no longer guaranteed to be convex and can give spurious or multiple solutions for the same data set – in the same spirit as a self-driving car miscategorizing a “STOP” sign as an indication to speed up or some other action.^48^ This would happen because gradient minimization got stuck in some spurious local minima or even a saddle point on the learning landscape. The slow modes thus derived would be spurious and using them as a biasing CV or RC would lead to incorrect and inefficient sampling. This could naturally lead one to derive misleading conclusions.

While the concerns stated above and the approach in this work to address them should be applicable to more general instances of AI application in molecular simulations, here we focus on the problem of enhanced sampling through MD-AI iterations. We report a new and computationally efficient algorithm designed to screen the spurious solutions obtained in AI-based methods. Our central hypothesis is that spurious AI solutions can be identified by tell-tale signatures in the associated dynamics, specifically through poor time-scale separation between slow and fast processes. Thus, different slow mode solutions obtained from different instances of AI applied to the same data set can be ranked on the basis of how much slower the slow mode is relative to the fast modes. This difference between slow and fast mode dynamics is known as spectral gap. We would like to emphasize that the concept of largest spectral gap correlating with CV optimality is a well-founded and theoretically justified concept at the heart of many previous studies.^49–53^ However, it has not yet been applied in a computationally tractable manner to representations arising from AI frameworks used on biased datasets, as is done in this work. Here this is made feasible through the use of the “Spectral Gap Optimization of Order Parameters (SGOOP)” framework.^54^ This builds a maximum caliber or path entropy^55^ based model of the unbiased dynamics along different AI based representations even when the underlying observables arise from biased simulations, which then yields spectral gaps along different slow modes obtained from AI trials. We demonstrate this path entropy based screening procedure in the context of our recent iterative AI-MD scheme “Reweighted Auotencoded Variational Bayes for enhanced sampling (RAVE)”.^40^ Here we show how this automated protocol can be applied to the study of a variety of molecular problems of increasing complexity. These include conformational dynamics in a model peptide, ligand unbinding from a protein, and extensive sampling of the folding/unfolding of the C-terminal domain of protein G (GB1-C16). We believe the presented algorithm marks a major step forward in the use of fully automated AI-enhanced MD for the study of complex bio-molecular processes.

### Theory

#### AI can mislead

In this work our starting point is the recent AI-based method RAVE.^37,40,56^ RAVE is an iterative MD-AI approach wherein rounds of MD for sampling are alternated with rounds of AI for learning slow modes. Specifically, RAVE begins with an initial unbiased MD trajectory comprising values of some order parameters **s** = (**s**_1_, **s**_2_, …, **s**_*d*_). These could be generic variables such as dihedrals or protein-ligand distances,^57^ as well as other CVs deemed to best describe the behavior of the system of interest. This trajectory is then treated with the past-future information bottleneck (PIB) framework.^58–62^ Per construction, the PIB is a low-dimensional representation with the best trade-off between minimal complexity and maximal predictive capability of the trajectory’s evolution slightly ahead in future. RAVE uses the PIB as a computationally tractable approximation for the RC which is traditionally considered as the definition of a slow mode.^63^ PIB is then used in an importance sampling framework to perform the next round of biased MD. Assuming that the biased PIB is close enough to the true slow mode or modes of the system, one expects the exploration of the configuration space in this new biased round of MD to be greater than in the previous round. The biased MD itself can be performed using one of the many available biased sampling schemes.^40,64,65^

In order to learn the PIB, RAVE uses an encoder-decoder framework. The PIB or RC *χ* is expressed as a linear combination of order parameters *χ* =Σ_*i*_ *c*_*i*_*s*_*i*_ where the order parameters are **s** = (**s**_1_, **s**_2_, …, **s**_*d*_), *c*_*i*_ denotes different weights^57^ and *d* denotes the dimension of the order parameter space. The PIB objective function which is then minimized in every training round can be written as a difference of two mutual informations: ^66^

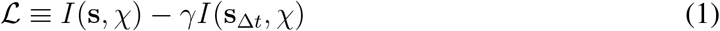

where *I*(.) denotes the mutual information between two random variables.^66^ The term *I*(**s**_Δ*t*_, *χ*) describes the predictive power of the model which is quantified by the amount of information shared by the information bottleneck *χ* and the future state of the system **s**_Δ*t*_ when the information bottleneck is decoded back to the order parameters space. To optimize the objective function, the information bottleneck *χ* should be as informative as possible about the future state of the system, quantified through increasing *I*(**s**_Δ*t*_, *χ*). At the same time, we seek to minimize the complexity of the low dimensional representation. Therefore, when the encoder maps the present state of the system **s** to information bottleneck *χ*, we aim to minimize the amount of information shared between them by decreasing *I*(**s**, *χ*). The parameter *γ* is introduced to tune the trade-off between predictive power and the complexity of the model.

In Eq. 1, the encoder is a linear combination of the input coordinates, thereby keeping it interpretable and relatively robust to overfitting. The decoder is a deep artificial neural network (ANN). Due to the principle of variational inference^40,67^ wherein optimizing the decoder is guaranteed to lead to a convex optimization problem, we are not concerned with over-fitting in the decoder. It is fitting the encoder, which directly leads to an interpretable RC, that is of concern to us here. This can be best illustrated through a simple numerical example involving protein conformational dynamics, which we describe in detail in **Methods** and in Fig. 1. We performed 6 different, independently initialized trials of PIB learning using the same input trajectory for a model peptide (alanine dipeptide), each running for same number of epoches. The RC was expressed as a linear combination of the sines and cosines of various Ramachandran dihedral angles. As can be seen in Fig. 1B, we obtain different RCs with different trials even though they are all stopped at the same low value of the loss function (within 4 decimal digits of precision). Given the use of an interpretable linear encoder, one can see a sense of symmetry even in the at first glance different looking RCs in Fig. 1B. However as we show later, the situation exemplified here exacerbates quickly with more complicated systems, and we expect this degeneracy to get only worse in more ambitious AI-based applications where even the encoder is non-linear,^37,41,51^ and/or where one does not really know *a priori* when to stop the training.

**Figure 1:**
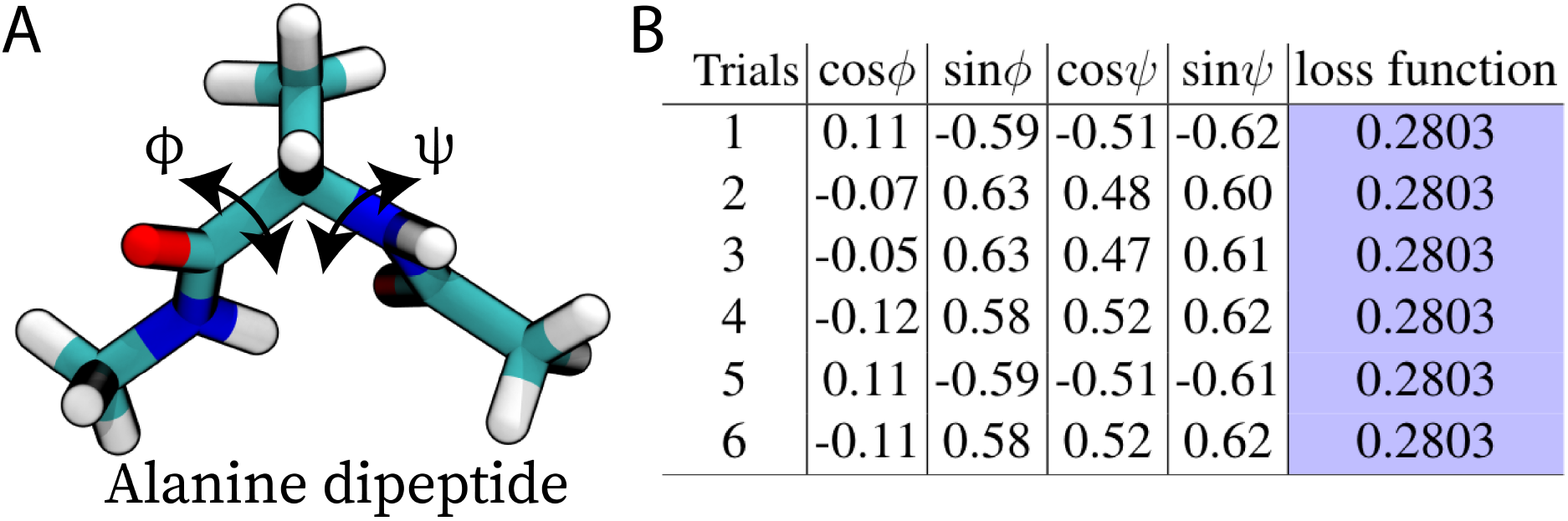
Spurious AI solutions for RCs describing conformational dynamics of alanine dipeptide. (A) Molecular representation of alanine dipeptide showing relevant Ramachandran dihedral angles, *ϕ* and *ψ*. (B) Table highlights the insensitivity of the objective function towards the changes in the weights of the order parameters. Six independently initiated trials of RAVE, on the same input trajectory, resulted in different RCs. The RCs are expressed as a linear combination of sines and cosines of *ϕ* and *ψ* with coefficients/weights listed in the table.

The above numerical example demonstrates the problem at heart of what we wish to tackle in this manuscript: how does one screen through spurious solutions resulting from attempts to optimize an objective function in AI applications to molecular simulations, and more broadly in chemistry and other physical sciences? The problem is especially difficult in two scenarios. Firstly, when one does not know the ground truth against which different AI solutions could be ranked, as is expected in any application where one seeks to gain new insight. Secondly, as is the case in AI-augmented MD, this problem will have critical, unquantifiable ramifications in iterative learning scenarios when any such AI-derived insight is used to make new decisions and drive new rounds of biased simulations. For instance in RAVE, we have yet another parameter that is not obvious how to select, namely the choice of the predictive time-delay Δ*t* in Eq. 1. As shown in Ref.,^68^ theoretically speaking the method is robust to the choice of this parameter as long as it non-zero yet small enough. In practice, it can be hard to judge whether it is indeed small enough or not.

### Path entropy model of dynamics can be used to screen AI solutions

In order to rank a set of AI-generated putative RCs, we appeal to the fundamental notion of time-scale separation, which is ubiquitous across physics and chemistry for example through concepts such as Born-Oppenheimer approximation^69^ and Michaelis-Menten principle.^70^ We posit that given a basket of RC solutions generated through AI, we can rank them as being closer to the true but unknown RC if they have a higher time-scale separation between slow and fast modes. Thus, a spurious AI solution should have a tell-tale signature in its dynamics, with poor separation between slow and fast modes. Indeed, one of the many definitions of an RC in chemistry is one that maximizes such a time-scale separation.^71^ To estimate this time-scale separation efficiently and rank a large number of putative AI based solutions for the true RC or PIB, here we use the SGOOP framework,^54^ which uses a maximum path entropy or caliber model^55,72^ to construct a minimal model of the dynamics along a given low-dimensional projection. To construct such a model, SGOOP requires two key inputs. First, it needs the stationary probability density along any putative RC, which we directly obtain after each round of RAVE.^40,68^ Second, it needs estimates of unbiased dynamical observables which we obtained from short MD simulations. With these two key sets of inputs, SGOOP can construct a matrix of transition rates along any putative RC. Diagonalizing this matrix gives the eigenvalues for the dynamical propagator. The spectral gap from these eigenvalues is then our estimate of the time-scale separation.^73^ While improving the quality of the dynamical observables can lead to increasingly accurate eigenvalues,^74^ here we use a computationally inexpensive dynamical observable denoted *(N)* and defined as the average number of nearest neighbor transitions per unit time along any RC. SGOOP protocol requires a standard grid parameter (also used for histogramming) which in all the studied systems was set to 20. We use *p*_*n*_ to denote the stationary probability density along any suitably discretized putative RC at grid index *n*. With these inputs, the SGOOP transition probability matrix *K* for moving between two grid points *m* and *n* is given by:^55,73^

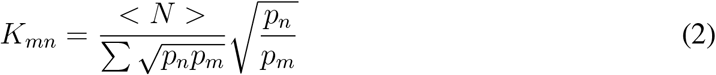

Our net product is an iterative framework that leverages the predictive power of RAVE and the fundamental notion of time-scale separation of SGOOP to generate an optimal RC. The use of AI in RAVE allows one to generate several possible candidate RCs, and by constructing a minimal path entropy based dynamical model we efficiently screen out spurious solutions generated from AI. We would like to note that maximum path entropy does not require any additional simulations beyond those already available from RAVE, rather its a post processing protocol which can be employed after each set of RAVE runs to sieve-out spurious solutions. The RC so identified is then used as a biasing variable in enhanced sampling, and the biased trajectory itself is fed back to the AI module to further optimize the RC. The iteration between this framework and sampling continues until multiple transitions between different intermediate states are achieved. We also apply this framework to cleanly select the best choice of predictive time-delay (Δ*t*) in Eq. 1 - the optimal predictive time-delay in our model for PIB is the one that achieves the highest time-scale separation.

## Results

In the previous section we described a path entropy and time-scale separation based paradigm to capture spurious solutions in AI-enhanced MD. In this section we illustrate the effectiveness of our framework through three generically relevant biophysical examples of increasing complexity. Specifically, we consider (A) conformational dynamics of a model peptide in vaccum, (B) dissociation of a millimolar-affinity ligand from FKBP protein, and (C) folding of the GB1-C16 peptide. All simulations are done at an all-atom resolution, including explicit water in (B) and (C). In all three systems, starting with an initial unbiased MD trajectory comprising of generic order parameters *s*, we perform iterative rounds of RAVE followed by biased enhanced sampling, using SGOOP to screen RC candidates generated in RAVE and to select the optimal time-delay Δ*t* in Eq. 1. Apart from the starting choice of order parameters which are kept quite generic (Table S1), all steps are carried out with minimal use of human intuition. To display the versatility of our framework, we combined it with two different enhanced sampling algorithms.^75^ In systems (A) and (B), we employ static biases to further enhance the conformational sampling of the model peptide and ligand dissociation along the reaction path.^76^ These static biases were directly obtained by inverting the probability distribution learnt during RAVE.^40,56^ In system (C), we employ time-dependent biasing through well-tempered metadynamics^24,65^ to capture folding of the GB1-C16 peptide. All the simulations were performed with GROMACS version 5.0^77^ patched with PLUMED version 2.4.2.^78,79^

### A. Conformational dynamics of alanine dipeptide

The first system we consider here is the well-studied case of alanine dipeptide in vacuum. It can exist in multiple conformations separated by barriers and commonly characterized by differing values of its backbone dihedral angles *ϕ* and *ψ* (Fig. 1B). Enabled by the small size of the system, we performed 3 independent simulations, each 2 *µ*s long. The corresponding trajectories along with the dihedral angles *ϕ* and *ψ* are provided as Supplementary Information (SI) in Fig. S1. In line with standard practice,^80,81^ the sines and cosines of these two dihedral angles provide natural input order parameters (OPs) *s* = (cos *ϕ*, sin *ϕ*, cos *ψ*, sin *ψ*) for RAVE, which then learns the optimal RC *χ* as a linear combination of these four. In the three independent trajectories, even with such long simulation times we capture only 1, 2, and 4 transitions between the axial and the equatorial conformations of the dipeptide. Using such input trajectories with different number of transitions helps us ascertain robustness of the protocol developed here. Each trajectory was used to perform RAVE with 11 different choices of the predictive time-delay Δ*t* in Eq. 1, ranging from 0 to 40 ps. Furthermore, 10 different trials were performed for each Δ*t* value corresponding to different input trajectories. This amounts to a total of 330 RAVE calculations, with 110 for each input trajectory. Each trial was stopped after the same training time, and the loss function value after the training as well as the RC so-obtained were recorded.

As hinted at in **Introduction**, we obtain very different RCs for the different Δ*t* values and for different independent trials. Furthermore, different trials that were stopped at similar loss function value gave different RCs and spectral gaps (Fig. 3 and S2). However our protocol of using spectral gaps to rank these different solutions works well in screening out the RC. In Figs. 3(A-C) we demonstrate the noisy correlation that we find between the loss function value and the spectral gap for all 3 input trajectories. In SI (Fig. S2B), we provide an illustrative figure for one particular trajectory showing how the same loss function value results in RCs with different free energy profiles, and that the one with the highest spectral gap stands out with most clearly demarcated metastable states. Similarly, the spectral gap captures the most optimal RC not just from the set of multiple trials at each time-delay, but it can also be used to select the optimal time-delay itself (Fig. 3D). In the subsequent calculations optimal time-delay of 8 ps, corresponding to the maximum spectral gap, was employed. Irrespective of the choice of input trajectory, we find that the optimal RC shows higher weights for *ϕ* (as compared to *ψ*) (Table 1), in line with previous studies which highlighted *ϕ* to be a more important degree of freedom than *ψ*.^40,82^ Using the RC corresponding to the *ntrans*= 4 and its probability distribution as a fixed bias,^40^ we then explored the conformational space of the peptide. The two-dimensional free-energy landscape along the dihedrals *ϕ* and *ψ* was able to capture axial and equatorial conformations of the peptide in only 20 ns of biased simulation (Fig. 3E). This is in excellent agreement with previously published studies for this system.^24,40^ However, biased simulations with the RAVE-alone RC result in poorer sampling of the configuration space relative to biased simulations using the RC further screened with SGOOP, as can be seen from the lesser number of transitions between the energy basins (Fig S3).

**Table 1:** Optimal weights of OPs obtained through a combination of RAVE and SGOOP.

| Trajectory | Transitions | $\cos\phi$ | $\sin\phi$ | $\cos\psi$ | $\sin\psi$ |
| --- | --- | --- | --- | --- | --- |
| 1 | 1 | -0.71 | -0.68 | 0.09 | -0.15 |
| 2 | 2 | 0.35 | 0.81 | 0.32 | 0.34 |
| 3 | 4 | 0.63 | -0.65 | 0.03 | 0.42 |

**Figure 2:**
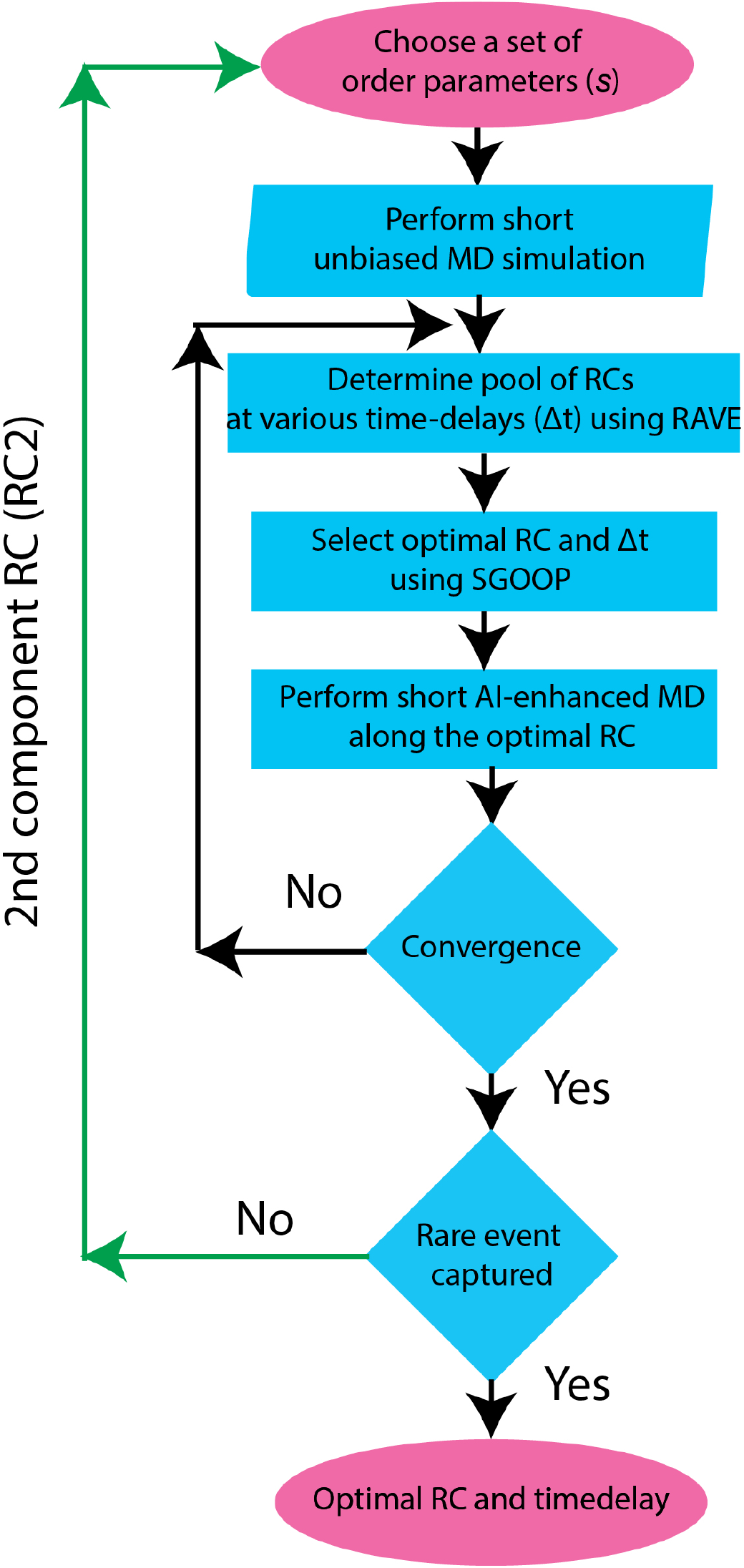
Flowchart illustrates our novel and computationally efficient protocol to screen AI solutions. Starting from short unbiased MD simulations, our protocol automatically screen the spurious solutions obtained in AI-based method and learns the optimal RC. In this work we demonstrate the applicability of our protocol in the context of RAVE and the screening of the spurious solutions is achieved by a path entropy based procedure.

**Figure 3:**
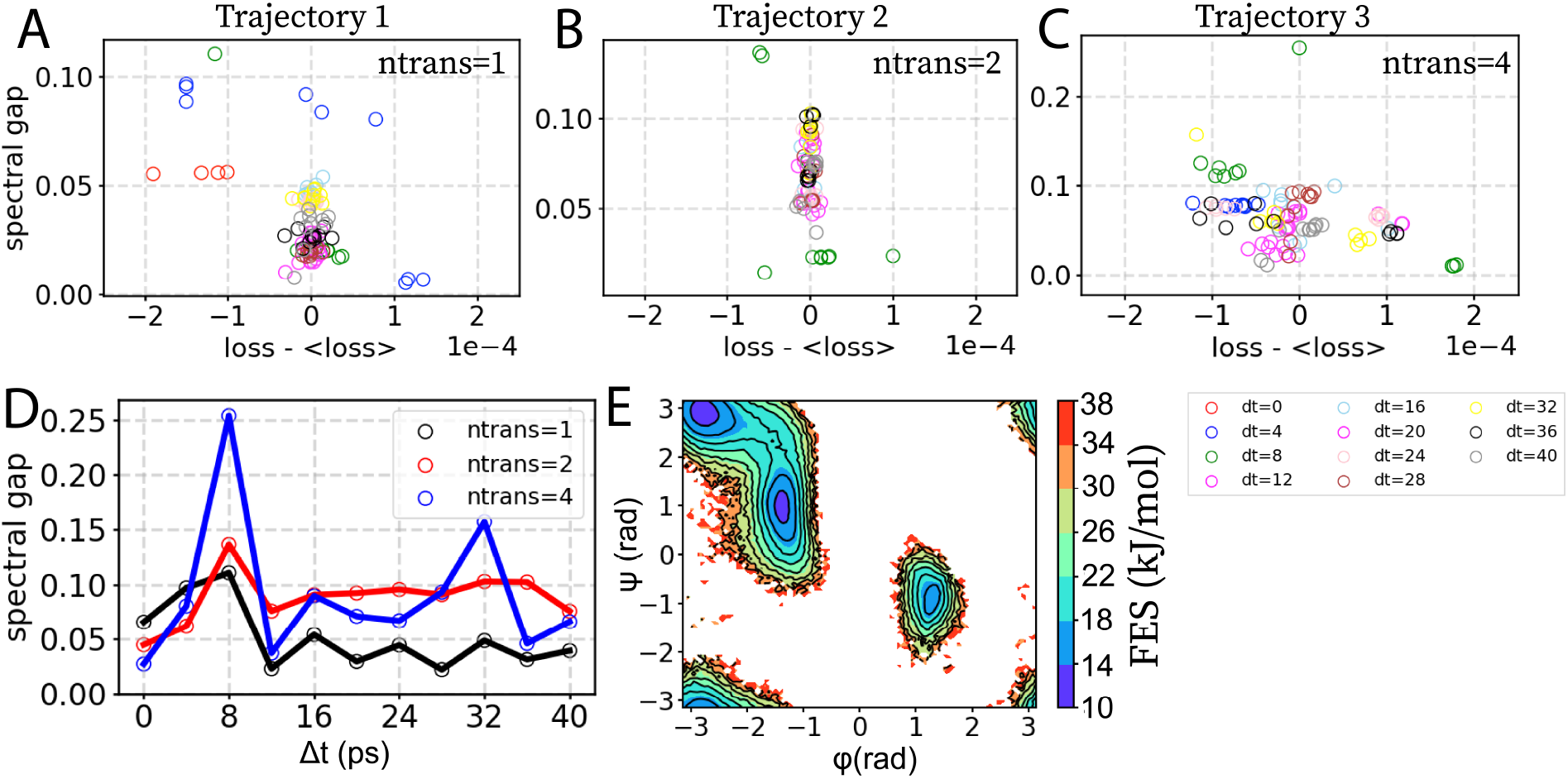
Capturing the spurious AI solutions in alanine dipeptide. Spectral gap and loss function values were calculated for each of the three unbiased trajectories at multiple time-delays Δ*t* between 0 and 40 ps, indicated using circles of different colors in the bottom right side of the figure. (A-C) show noisy correlation between the loss and the spectral gap for number of transitions *ntrans* equaling 1, 2, and 4 respectively. Different circles denote different independent trials, with color denoting Δ*t*. For visual clarity, for every *ntrans* we have plotted a mean-free version of the loss function value by subtracting out the average of all losses. (D) Maximum spectral gap (out of 10 different trials of RAVE) vs. Δ*t* was plotted for 3 different unbiased trajectories. Optimal time-delay of 8 ps was employed in subsequent calculations. (E) Free energy surface (FES) along the two dihedrals Φ and Ψ, obtained from 20 ns-long simulation in the presence of static bias. Energy contours are shown at every 4 kJ/mol.

### B. Unbinding of millimolar-affinity ligand from FKBP

In the second example, we applied our framework to a well studied problem of dissociation of 4-hydroxy-2-butanone (BUT), a millimolar affinity ligand, from the FKBP protein (Fig. 4A). Force-field parametrization^83–85^ and other MD details are provided in SI. Here, our objective was to use RAVE to learn the most optimal RC on-the-fly as well as the absolute binding free energy of this protein-ligand complex. This is a difficult and important problem for which many useful methods have been already employed with varying levels of success.^86^ At this stage at least, our intention is not to compete with these other existing methods, but instead validate that our framework works for a well studied benchmark problem. We begin by performing four independent MD simulations of FKBP in its ligand-bound form (PDB:1D7J).^87^ The MD simulations were stopped when the ligand unbound, specifically when it was 2 Å away from the binding pocket (Fig. 4B). All trajectories were expressed in terms of 8 OPs representing various distances between the center of mass (COM) of the ligand and the COM of the residues in the binding pocket (Table S1), which comprise a natural choice for the process of ligand unbinding from the protein and have been employed in previous studies.^57,88^ We combined the results of the four independent MD trajectories to perform RAVE with 11 different choices of predictive time-delay, ranging from 0 to 40 ps. At each Δ*t*, 10 different trials were performed resulting in a total of 110 RAVE calculations. Each trial was stopped after the same training time, and the loss function value after the training as well as the RC so-obtained were recorded. Different RCs were screened using a path entropy based model as discussed in **Theory** and done for alanine dipeptide. We again find noisy correlation between the loss function values and the spectral gap (Fig. 4C), for the case of Δ*t* = 40 ps (additional plots are given in Fig. S4). The same value of loss function gives rise to very different values of the spectral gap and of the RC (Fig. 4C). Furthermore, the spectral gap not only captures the most optimal RC, it is also able to select the most optimal time-delay (Fig 4D). By using this RC and its probability distribution as a fixed bias,^40^ we then performed 800 ns of biased simulations starting from the bound pose, but allowing the ligand to re-associate (Fig. S5A,B). Through this we then calculate the absolute binding affinity of the protein-ligand complex to be 6.6 kJ/mol (Fig 4E), in good agreement with values reported through metadynamics.^57^Interestingly, the binding affinity of the protein-ligand complex was also in good agreement with the values reported through extended unbiased simulations by Pan et al.,^89^ although, the ligand was parameterized with the generalized amber force field (GAFF).^88^ It is worth pointing out that the ANTON simulations took 39 *µ*s while we obtained converged estimates in around 800 ns, reflecting roughly a factor of 48 speed-up with minimal use of prior human intuition.

**Figure 4:**
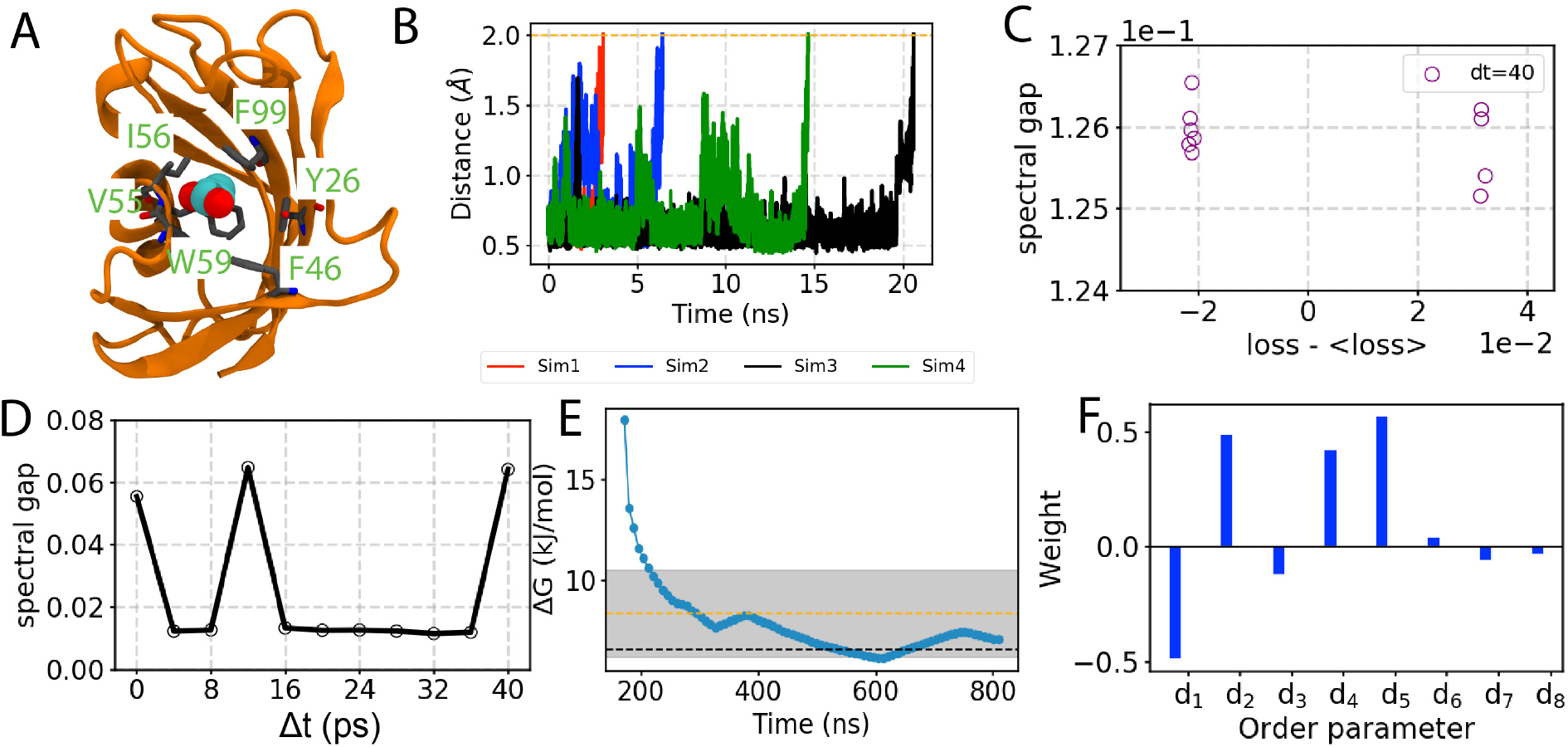
Unbinding of 4-hydroxy-2-butanone (BUT) from FKBP. (A) Molecular image of the bound FKBP/BUT protein–ligand complex, with binding pocket residues highlighted. The distances to the residues were used as different OPs detailed in Table S1. (B) Time evolution of the distance between the center of mass (COM) of the bound ligand and COM of residue W59. It is to be noted that in order to avoid entropy dominant process, only the ligand-bound trajectories were considered in our protocol. (C) Spectral gap and loss (at time-delay of 40 ps) for 10 different trials were calculated after combining all four independent trajectories at multiple time-delay Δ*t* between 0 and 40 ps, indicated using circles. Rest of the time-delays are shown in Fig. S1. For visual clarity, at each iteration, we have plotted a mean-free version of the loss function value by subtracting out the average of all losses. (D) Plot of maximum spectral gap (out of 10 different trials of RAVE) vs time-delay (Δ*t*). (E) Absolute binding free energy G in kJ/mol of FKBP/BUT system as a function of simulation time with static external bias. The dotted black and orange line shows the reference value reported through metadynamics^57^ and long unbiased MD simulations performed on ANTON.^89^ The shaded region shows the free energy estimate from long unbiased MD simulations performed on Anton including the *±*2.092 kJ error reported.^89^ (F) A visual depiction of the OP weights.

The use of a linear encoder in RAVE allows us to directly interpret the weights of the different OPs in the RC (Fig. 4F). The highest weight corresponds to the OP *d*_5_, which is the ligand separation from residue I56. This residue forms direct interactions with the bound ligand in the crystal structure. Interestingly, previous studies^87^ have highlighted the importance of I56 as it forms hydrogen bonding interactions with the carbonyl group of the bound ligand, our algorithm also captured it as the most significant OP. Followed by this highest weight component, the second and third highest components are for *d*_1_ and *d*_2_, denoting respectively distances from the residues V55 and W59. These are roughly equal in magnitude, reflecting that the ligand moves closer to V55 and W59 as it moved away from I56.

### C. Folding/unfolding dynamics of GB1 peptide

Finally we tested our method on the folding/unfolding dynamics of GB1-C16, which is known to adopt a *β*-hairpin structure.^90–94^ Force-field parametrization and other MD details are provided in the SI. The free-energy landscape of this peptide has been extensively explored by replica-exchange MD simulations and clustering based methods.^90,94,95^ These studies reported the presence of multiple intermediate conformations by projecting the simulation data along multiple OPs, such as radius of gyration (Rg), root-mean-squared deviation (RMSD), fraction of native contacts (NC), and native state hydrogen bonds (NHB). These OPs on their own were not able to distinguish between intermediate conformations with proper energy barriers. However, using a combination of these OPs as input in advanced slow mode analysis methods such as TICA^52,96,97^ recovers a more superior two-dimensional description.^90^ That work however used more than 12 *µ*s of enhanced sampling, specifically replica exchange MD trajectories, for this purpose. Here instead we use just 1.6 *µ*s of unbiased trajectories as our starting point. From this point onwards, using the same OPs as in Ref.,^90^ our work provides a semi-automated solution in deriving an optimal two-dimensional RC for GB1-C16, which is capable of resolving the intermediate conformations. Here, in contrast to the previous two examples we use well-tempered metadynamics^24,65^ simulations as the choice of enhanced sampling engine coupled with RAVE.

We start by performing four independent 400-ns of unbiased MD simulations of the peptide in explicit solvent. All the simulated systems were observed to be fairly stable when projected along a library of OPs comprising RMSD, NC, Rg, and NHB, with their detailed construction described in SI (Fig. 5, Fig. S6). All the unbiased trajectories were mixed and fed into RAVE for learning the RC. We performed 10 different trials of RAVE for different time-delays Δ*t*, ranging from 0 to 20 ps which amounts to a total of 110 RAVE calculations. Different putative RCs learnt from RAVE were screened using the path entropy based model as discussed in the **Theory** and as done for the other two systems. Similar to the previous systems, we find noisy correlation between the loss function value and the spectral gap (Fig. S7). The most optimal RC was selected for biased simulations using well-tempered metadynamics (Fig. S7A,B). Based on the maximum spectral gap, we chose Δ*t*=8 ps for the next round of 50 ns-long metadynamics simulation. We then alternatively iterate between the rounds of learning improved RC, using our framework, and running metadynamics using the optimal RC in every iteration. After two iterations, we did not find any further improvement in sampling with this one-dimensional RC, which we call *χ*_1_. With a 1-d metadynamics we were unable to attain back and forth transitions between different metastable states, suggesting the presence of missing/orthogonal degrees of freedom not encapsulated by *χ*_1_. In order to learn these other degrees of freedom through the second component of the RC, which we call *χ*_2_, we used the protocol from Ref.^98^ For practical purposes, this corresponds to ignoring the already learnt *χ*_1_ and treating the biased trajectory without any consideration of the bias along *χ*_1_. We would like to note that in the previous study^73^ we have extended the scope of SGOOP by employing a the notion of conditional probability factorization where known features are effectively washed out to learn additional features of the underlying energy landscape. This is what we have used for RAVE as well in the current work. In principle RAVE could be directly used to output a two-dimensional or even higher-dimensional RC, but this protocol ensures that we gradually ensure the RC dimensionality only when a lower dimension is found insufficient for sampling. We then performed 50 ns long 2D metadynamics simulations (Fig. S7 E,F), which were used to train *χ*_2_. The most optimal 2-dimensional RC obtained after three iterations of training *χ*_2_ is detailed in (Fig. 6A). The backbone heavy atom RMSD contribute the most toward the construction of slowest dimension *χ*_1_, whereas Rg contributed more towards the second slowest dimension.

**Figure 5:**
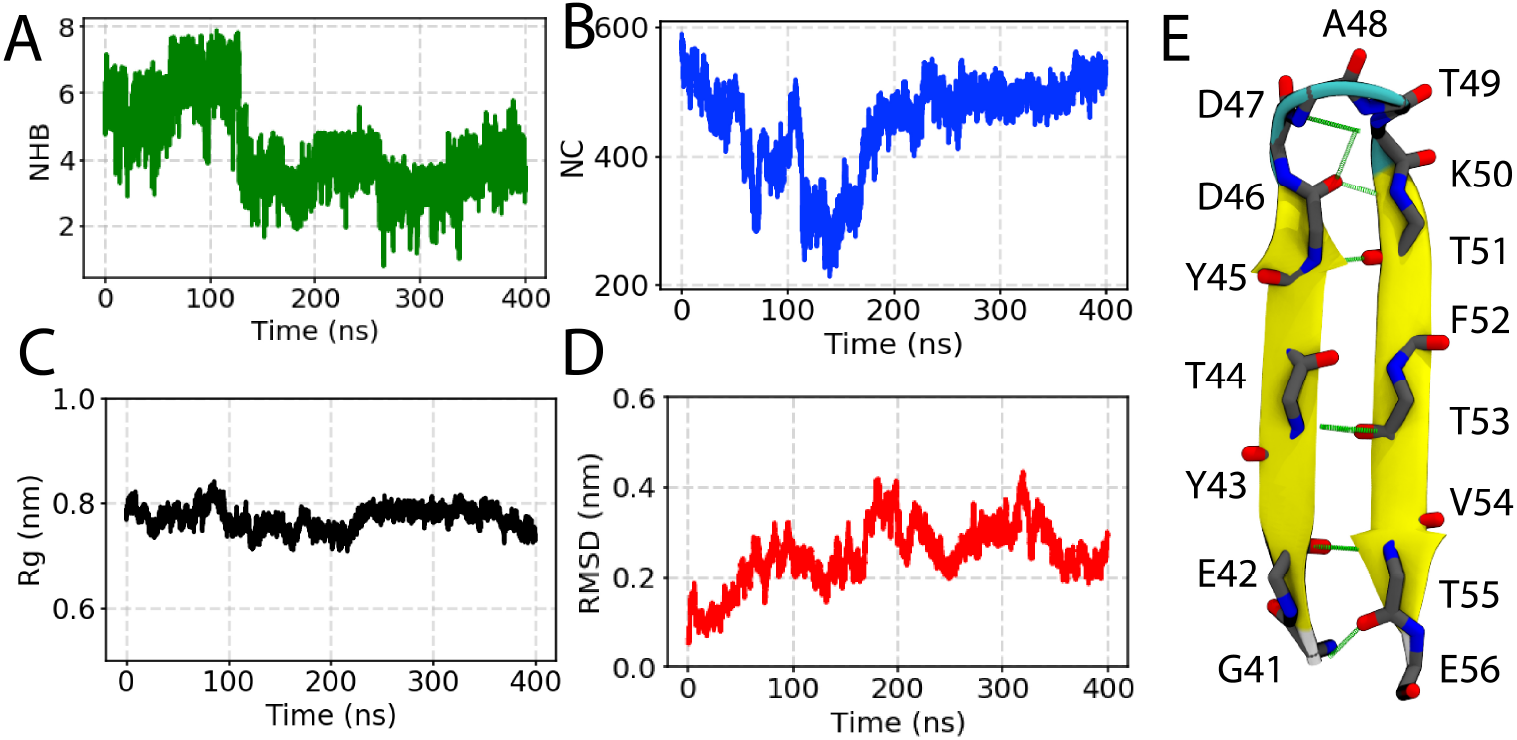
Dynamics of GB1-C16 captured from unbiased MD. One of the four representative trajectory of the peptide in explicit solvent is projected along different order parameters (A) number of hydrogen bonds (NHB), (B) native contacts (NC), (C) radius of gyration (Rg), and (D) root-mean square displacement (RMSD). (E) Molecular image of the GB1-C16. Native backbone hydrogen bonds are highlighted with green lines.

**Figure 6:**
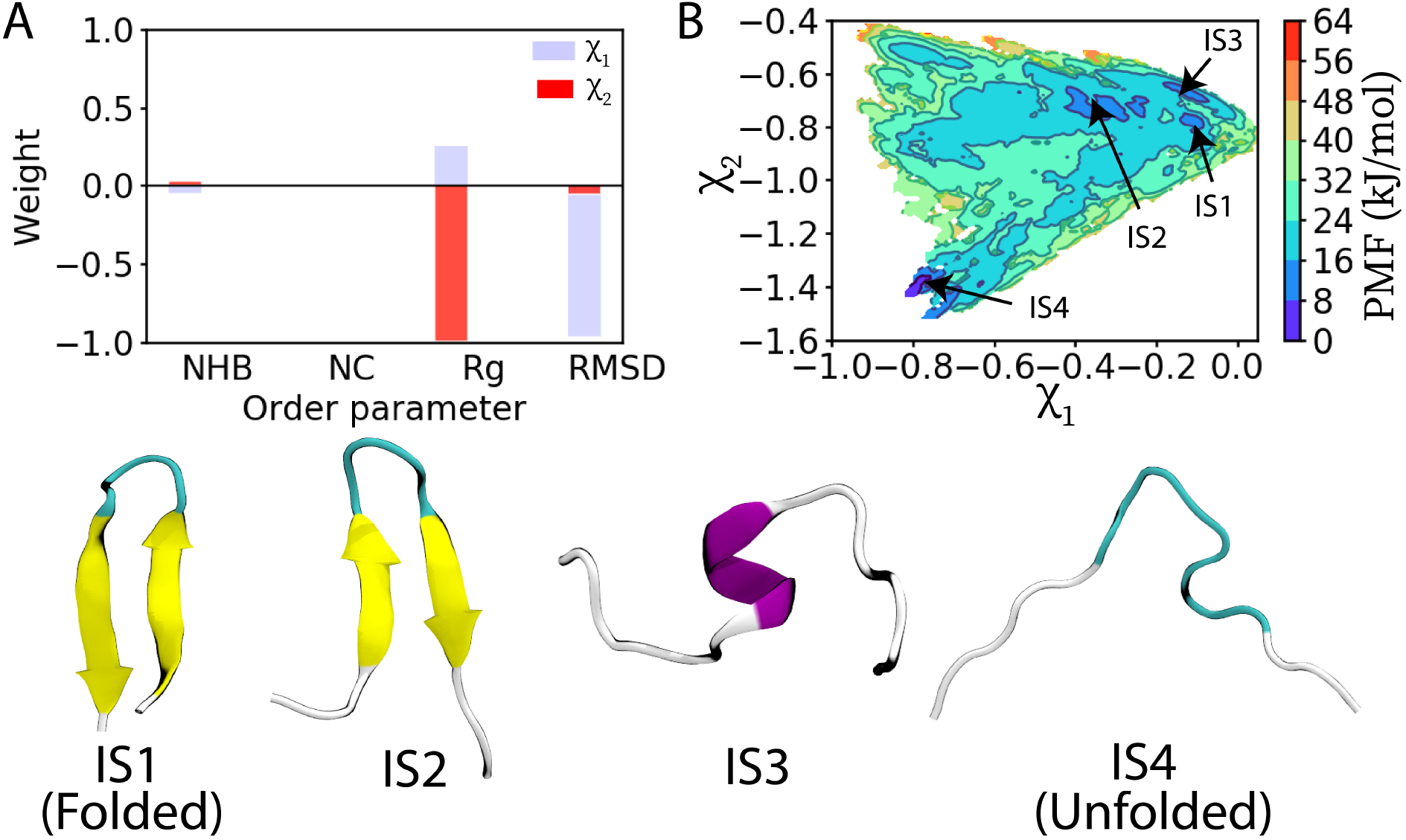
Free-energy landscape and OPs contribution. (A) Contribution of the different Ops to the 2-dimensional RC *χ*. The two components *χ*_1_ and *χ*_2_ are shown in blue and red bars, respectively. (B) A highly rugged 2-dimensional free-energy landscape of GB1-C16 folding/unfolding. We were able to capture multiple states, corresponding to the folded (IS1), unfolded (IS4), and intermediate states (IS2 and IS3). Interestingly, it is only by projecting the free energy as a function of the 2 RCs that we were able to capture a partially helical state (IS3), which otherwise was not easy to distinguish solely using traditional OPs. Representative snapshots of the captured structures are shown in the bottom panel, and their locations on the energy landscape shown in B.

The two-dimensional RC is then used in longer well-tempered metadynamics simulations to facilitate movement between different metastable states (SI Video 1) and to obtain converged free energy surfaces. We performed 1.2 *µ*s-long metadynamics simulations at 300 K, starting from the crystal structure (Fig. S8). The 2-dimensional metadynamics simulations were performed with initial hill height 0.5 kJ, bias factor = 10, Gaussian widths 0.03 for both *χ*_1_ and *χ*_2_, and bias added every 4 ps. Additional restraint potential was applied along the RMSD order parameter preventing very high values from being attained (see details in SI). In principle this step is not necessary as the simulation would eventually return back to low RMSD states, but in practice, due to the entropic nature of the high RMSD states, such a restraint significantly helps with computational efficiency. Fig. 6B shows the 2D free energy landscape as a function of the two RC components, at 300 K. We find that the system shows multiple energy basins corresponding to the different stable and metastable intermediates. Interestingly, we captured a helical conformation of this peptide, which was not easy to distinguish by using a combination of conventional OPs like RMSD, Rg, contact map, etc.^99^ For example, previous metadynamics-based studies employed Rg and native hydrogen bonds to accelerate the folding process, but they were not able to clearly demarcate distinct conformational states with energy-barriers.^100,101^ Interestingly,when the two-dimensional free-energy landscapes when projected along the pair of OPs, yields results consistent with the previous studies and suggests the presence of two metastable states (Fig. S9).

## Conclusion

To conclude, we have introduced a new approach to sieve out the spurious solutions from AI-augmented enhanced sampling simulations.^37,38^ AI-based approaches have had indisputable impact across sciences, including their use in enhancing the efficiency of molecular simulations. However, when these AI-based approaches are applied to a data sparse regime it can lead to spurious or multiple solutions. This would happen because gradient minimization can get stuck in some spurious local minima or even saddle points on the learning landscape, leading to misleading use of AI.

To deal with this issue of trustworthiness of AI in molecular simulations, we report a novel, automated algorithm aided by ideas from statistical physics.^102^ Our algorithm is based on the simple but powerful notion that a more reliable AI solution will be one that maximizes the time-scale separation between slow and fast processes. This fundamental notion of time-scale separation was implemented on the basis of maximum caliber- or path-entropy-based method, SGOOP.^54,55^ We would like to emphasize that our approach, and spectral gap based optimization in general^54^ might have as of yet unexplored connections with the Variational Approach for Markov processes (VAMP) .^103^ The framework developed here should be applicable to many recent methods (Ref.^37^ and references therein) which involve iterating between MD and AI for sampling and learning respectively. Here we demonstrate its usefulness through our recent integrated AI-MD algorithm RAVE.^40^ We illustrate the applicability of our algorithm through three illustrative examples, including the complex problem of capturing the energetic landscape of GB1 peptide folding in all-atom simulations. In this last case, we started from a library of 4 order parameters that are generic for folding/unfolding processes and demonstrated how to semi-automatically learn a 2-dimensional RC, which we then used in well-tempered metadynamics protocol to obtain folding/unfolding trajectories. This directly allows us to gain atomic level insights into different metastable states relevant to the folding/unfolding process. We thus believe that our method marks a useful and much needed step forward in increasing the utility of machine learning and AI-based methods in the context of enhanced sampling and one can expect that such an approach could be applicable to molecular simulations in general, although this is purely speculative at this point and remains to be verified.

## Acknowledgements

We thank the Donors of the American Chemical Society Petroleum Research Fund (No. PRF60512-DNI6) (PT), National Institutes of Health under award number P41-GM104601 (ET), the NCI-UMD Partnership for Integrative Cancer Research (YW), the COMBINE fellowship (DGE-1632976) (ZS), and Beckman Institute Graduate Fellowship (SP) for financial support. This work used the Extreme Science and Engineering Discovery Environment (XSEDE) Bridges through allocation CHE180027P, which is supported by National Science Foundation grant number ACI-1548562 (PT). We also thank UMD’s Deepthought2 and MARCC’s Bluecrab HPC clusters for computing resources. We would like to thank Navjeet Ahalawat and Jagannath Mondal for help with setting up the GB1 peptide system for MD simulation. We would also like to thank Andrew Ferguson for useful discussion and Joáo Marcelo Lamim Ribeiro for the careful reading of the manuscript.

## Data Availability

The data that support the findings of this study are available from the corresponding author upon reasonable request RAVE code is available at https://github.com/tiwarylab/RAVE and SGOOP code can be accessed at https://github.com/tiwarylab/SGOOP

## Supporting Information

### Methods

#### Molecular dynamics simulations

All the systems were simulated with GROMACS version 5.0^77^ patched with PLUMED version 2.4.2. ^78,79^ We constrained the bonds involving hydrogen atoms using LINCS algorithm and employed an integraion timestep of 2 fs. More details on individual systems and simulation protocols are provided below.

A. **Simulation setup for alanine dipeptide in vacuum** We followed Ref. ^40^ to set up our simulations for alanine dipeptide in vacuum. AMBER03 force field was used for the dipeptide. Three independent 2 *µ*s-long MD simulations each with randomized initial velocities were performed to improve the sampling. The temperature was kept constant at 300 K using the velocity rescaling thermostat. ^104^
B. **Simulation setup for the FKBP-BUT complex**. As a second system, we studied the dissociation of 4-hydroxy-2-butanone (BUT) from the protein FKBP. AMBER99SB-ILDN force field ^83–85^ was used for the protein along with TIP3P water, and the ligand was parametrized with the generalized amber force field (GAFF). ^105^ The simulations were initiated from the BUT-bound crystal structure of FKBP (PDB ID: 1D7J). Initial equilibration was performed in NPT ensemble for 1 ns followed by four independent production runs with Nosé-Hoover thermostat ^106^ maintaining the temperature at 300 K.
C. **Simulation setup for GB1 peptide**. As a third system, we studied the folding dynamics of GB1 peptide in water. The system was setup and equilibrated following the protocol described in Ref . ^90^ After this we performed four independent production runs, each lasting for 400 ns.

#### Metadynamics

The central idea behind metadynamics is that by adding a history-dependent biasing potential it encourages the given system to escape the free energy minima, and explores various stable and metastable regions of the energy landscape. ^24^ Following the reweighting procedure, one can then construct the underlying energy landscape along any RC. ^107^ To capture the unfolding energy landscape of GB1 peptide, we employed well-tempered metadynamics using PLUMED implementation. ^24,78,79^ The simulations were performed along the optimal RC obtained through the combination of RAVE and SGOOP. The metadynamics simulations were performed with the biasfactor of 10, initial hill height of 0.5 kJ/mol, and bias deposition rate of 4 ps. The sigma values of the added Gaussian bias was set to the standard deviation of the biasing RC obtained from the unbiased simulation 400 ns.

Once a 50 ns metadynamics run was completed, the biased trajectory was reweighted and used as an input for the second round of RAVE. ^40^ The system in this 50 ns biased runs explored more conformational space than in initial 400 ns of unbiased simulations. After this, we performed another round of biased simulations, where we employed 2D metadynamics along the optimal RCs obtained through the combination of RAVE and SGOOP. To prevent the peptide from getting trapped in entropic non-physical conformations due to the accumulated bias, we also restrain the movement of the peptide along the RMSD order parameter by addition of another bias. This bias had the form of Eq. 3. The final round of 2-dimensional metadynamics simulations were performed for 1200 ns.

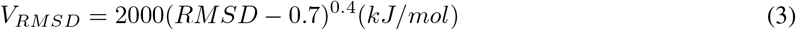

#### Definitions of Order Parameters

For each system we first constructed a dictionary of order parameters (OPs), summarized in Table 2.

**Supplementary Table S1:**
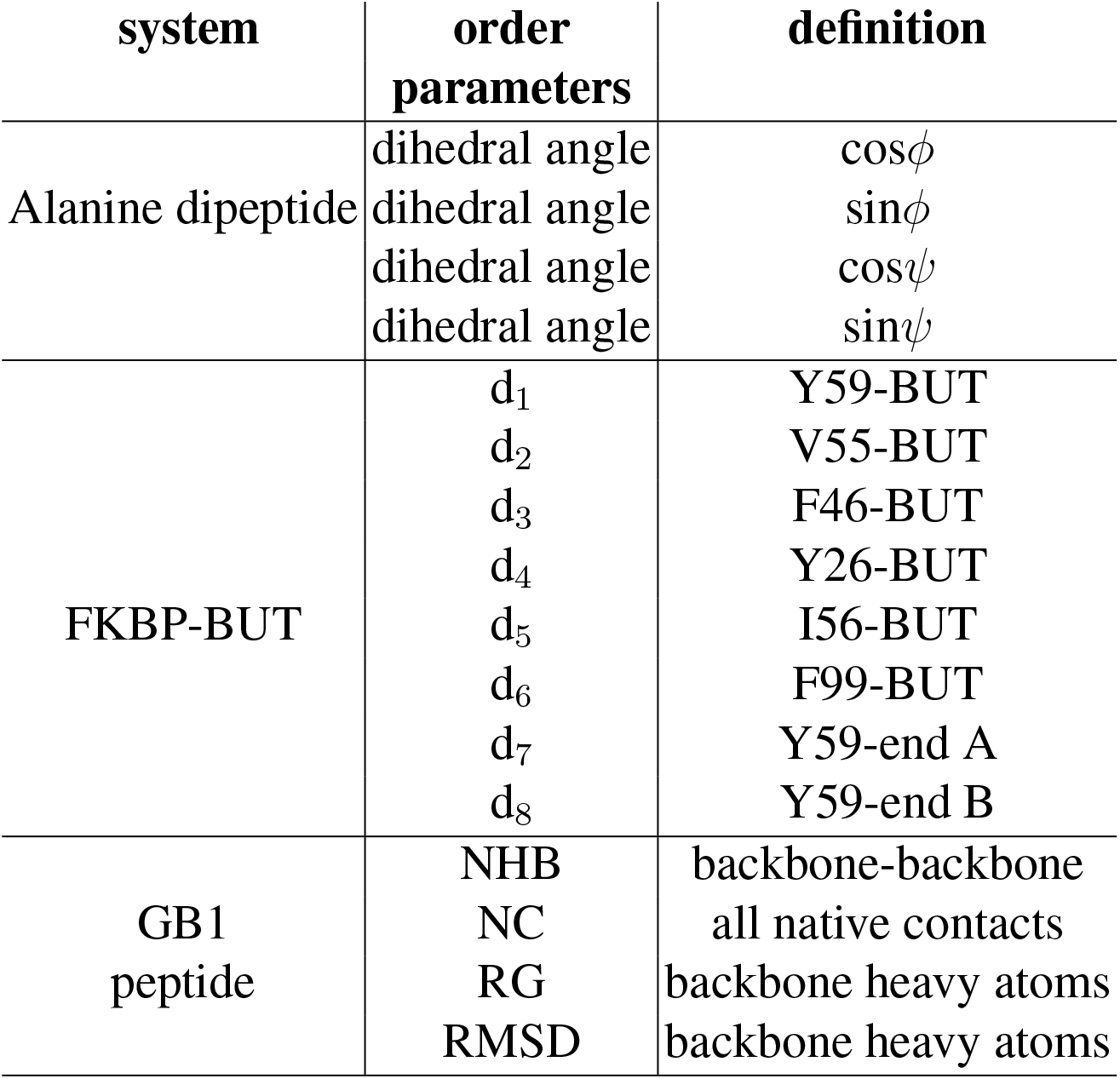
Details of the Pre-Selected Dictionary of Order Parameters (OPs)

redAll the distances in FKBP system was calculated with respect to the center of mass of the residues and the ligand. We have calculated NC and NHB using a switching function

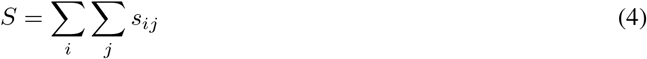

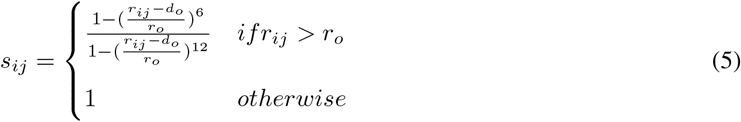

where d_*o*_= 0.05 nm and r_*o*_ = 0.65 nm were used for NC calculations and d_*o*_= 0.0 nm and r_*o*_ = 0.25 nm were used for NHB calculations. In the case of NC contacts, r_*ij*_ is a distance between *i* and *j* heavy atoms in the native state. While, in NHB, *i* and *j* are the pairs of O and HN (amino group) atoms of the backbone. RMSD of all the backbone heavy atoms with respect to crystal structure and RG of all the backbone heavy atoms were also calculated using PLUMED.

#### Neural network architecture

Hyper-parameters in this work included the variance of Gaussian noise, the number of neurons in hidden layers, initializer of weights of each layer, and the learning rate for the RMSprop algorithm. In the studied examples, all these hyper-parameters are set to be the same. The variance of Gaussians was kept 0.005 Each hidden layer had 128 neurons. The learning rate was set to be 0.003. Initial weights of each layer were randomly picked from a uniform distribution within range [0.005, 005]. We note that the independent RAVE trials even when used on the same trajectory are randomized, because

1. We randomly initialized the parameters in neural network.
2. Randomly generated seed was used to shuffle the training data.

**Figure S1:**
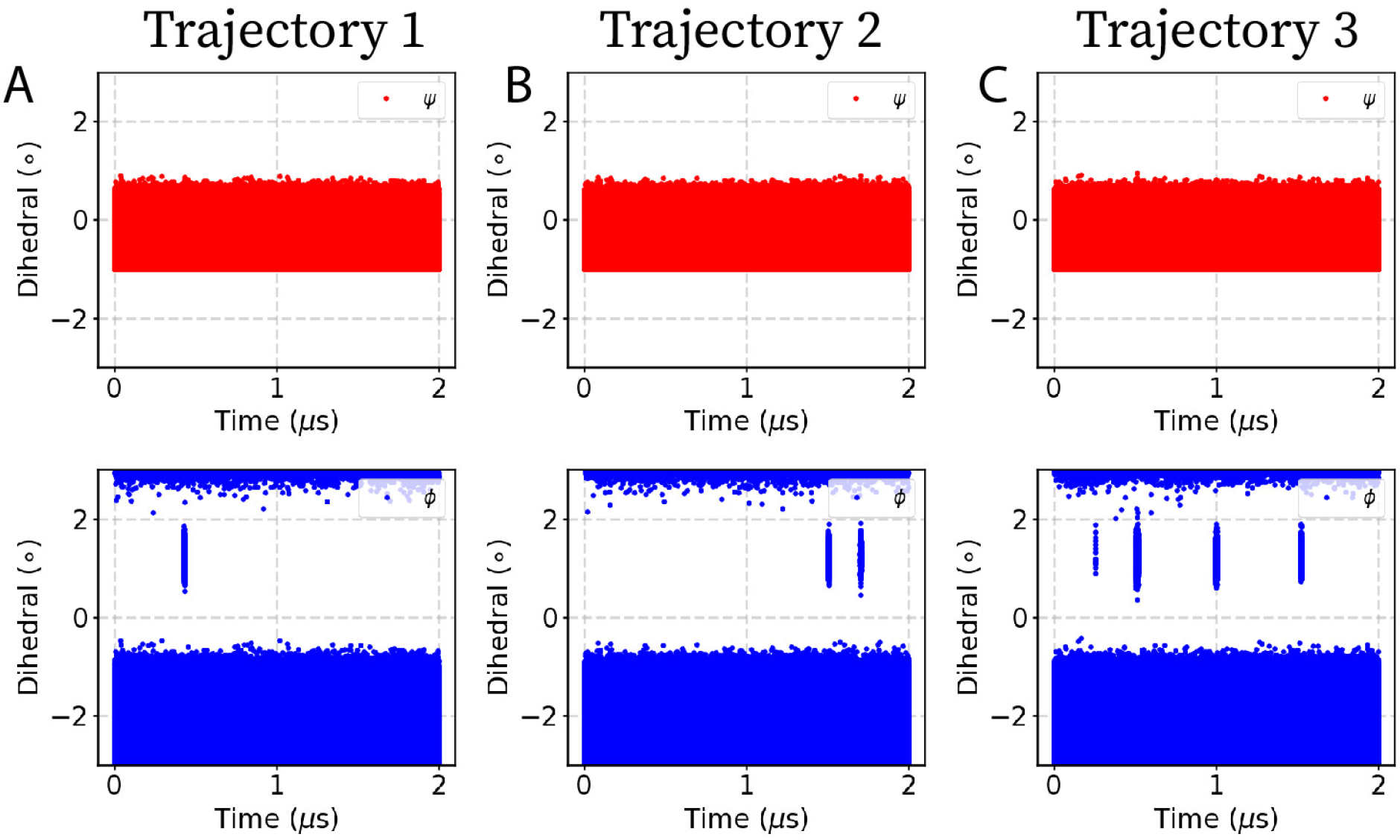
Unbiased MD simulations of model peptide in vacuum. (A-B) Three independent MD simulations of alanine dipeptide projected along dihedral (*ϕ, ψ*) angle space. Differing number of transitions along the *ϕ* order parameter were captured in the different independent simulations.

**Figure S2:**
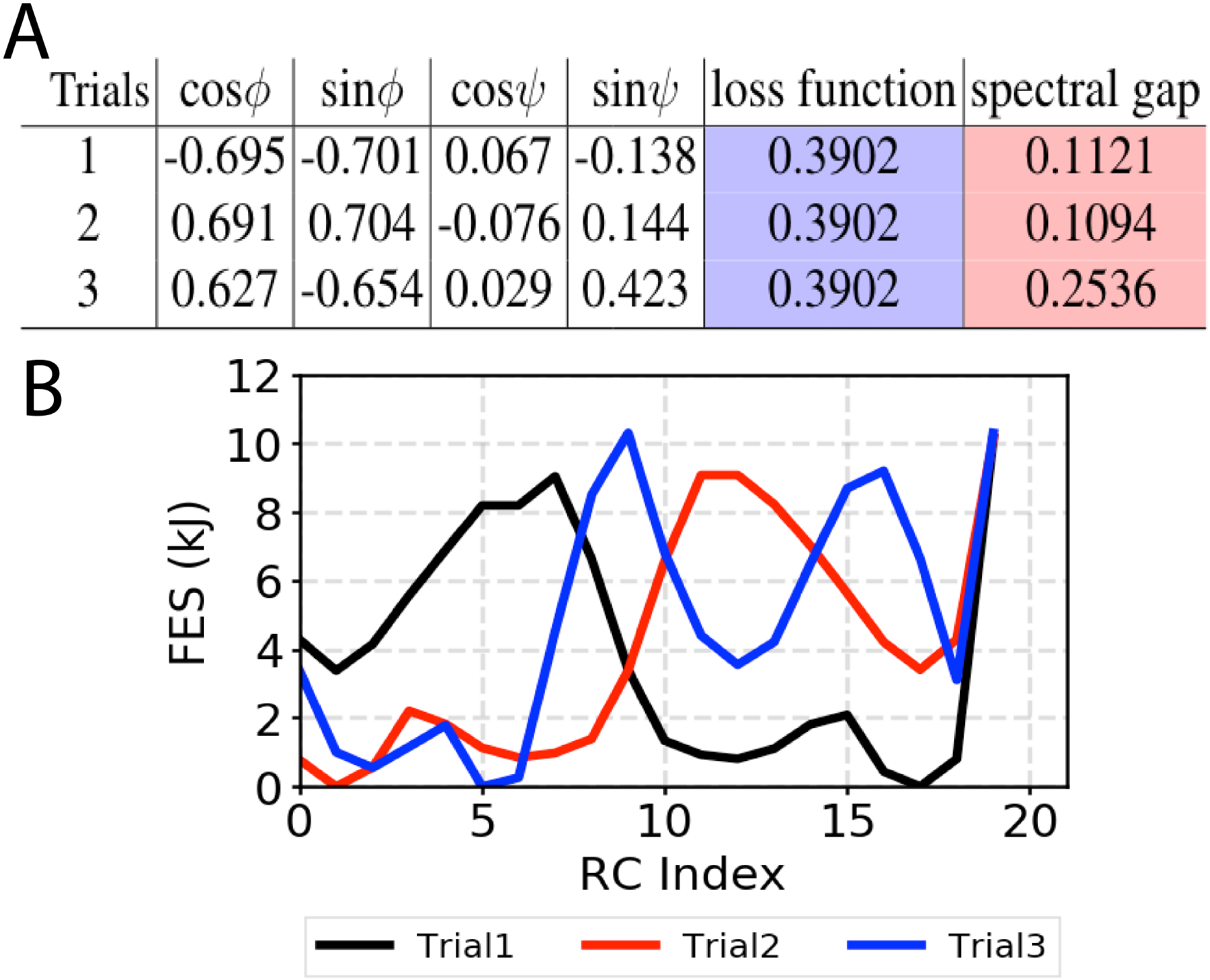
Same loss function can give differing RCs. (A) Table highlights the sensitivity of spectral gap towards changes in the weights of the order parameters. While objective function remain insensitive to the changes in the weights of the order parameter. (B) Free energy landscape of one particular trajectory showing same loss function results in RCs with very different free energy profiles. The RC with maximum spectral gap highlights distinct metastable state. The different local minima in this plot correspond to different metastable states which are not marked here to avoid confusion, and as our main aim is to illustrate how the free energies along different RCs from RAVE might appear different.

**Figure S3:**
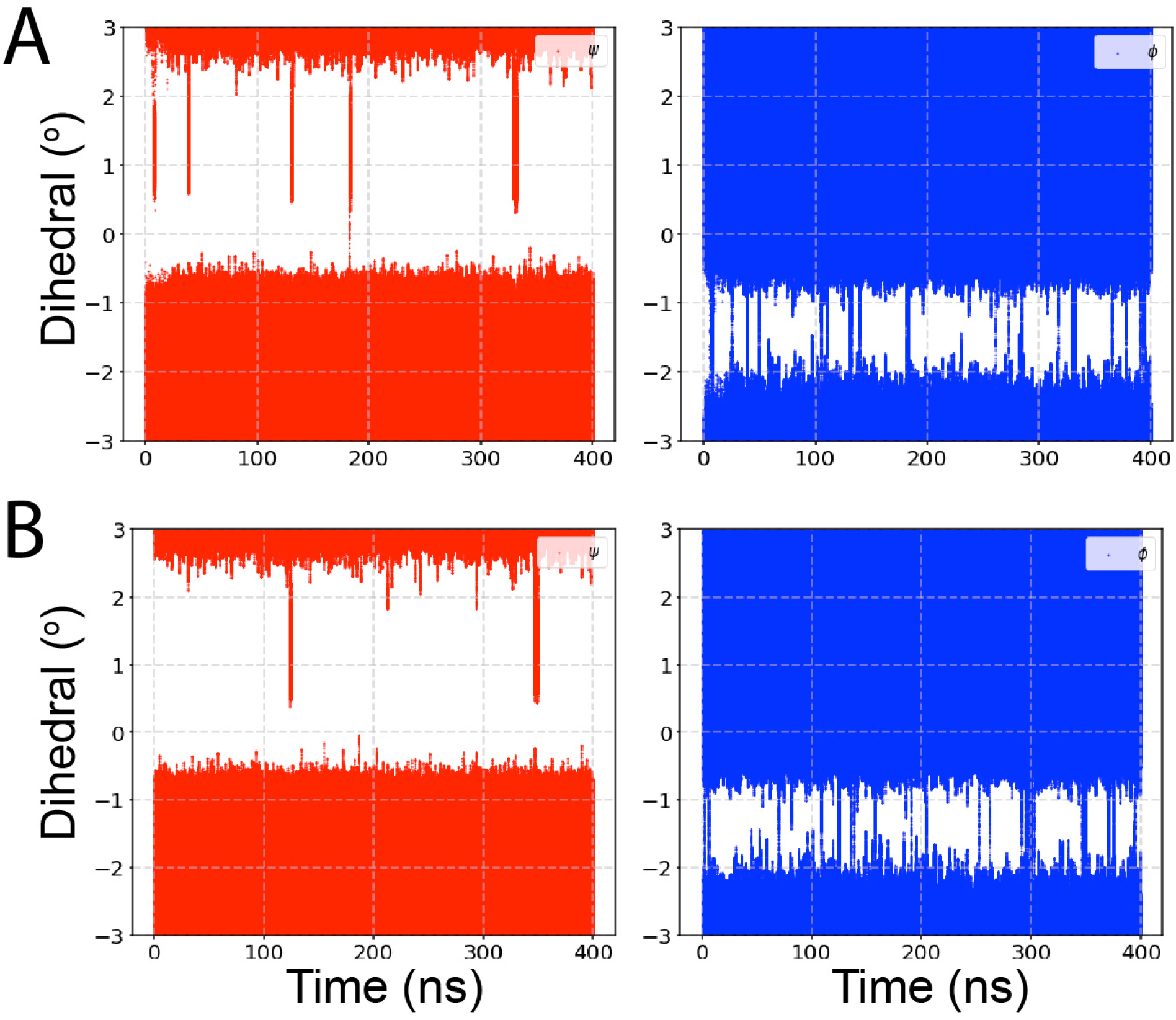
Time evolution of *ϕ* and *ψ* angles after one round of RAVE. Static biased MD simulations of alanine dipeptide along (A) optimal RC screened with SGOOP as having high spectral gap, and (B) less optimal RC without SGOOP screening. The precise RC definitions are provided in the Results section in the main text. Trajectories are shown for dihedral (*ϕ, ψ*) angle space. We observed more transitions when biased along the optimal RC, while both being relatively superior to unbiased MD.

**Figure S4:**
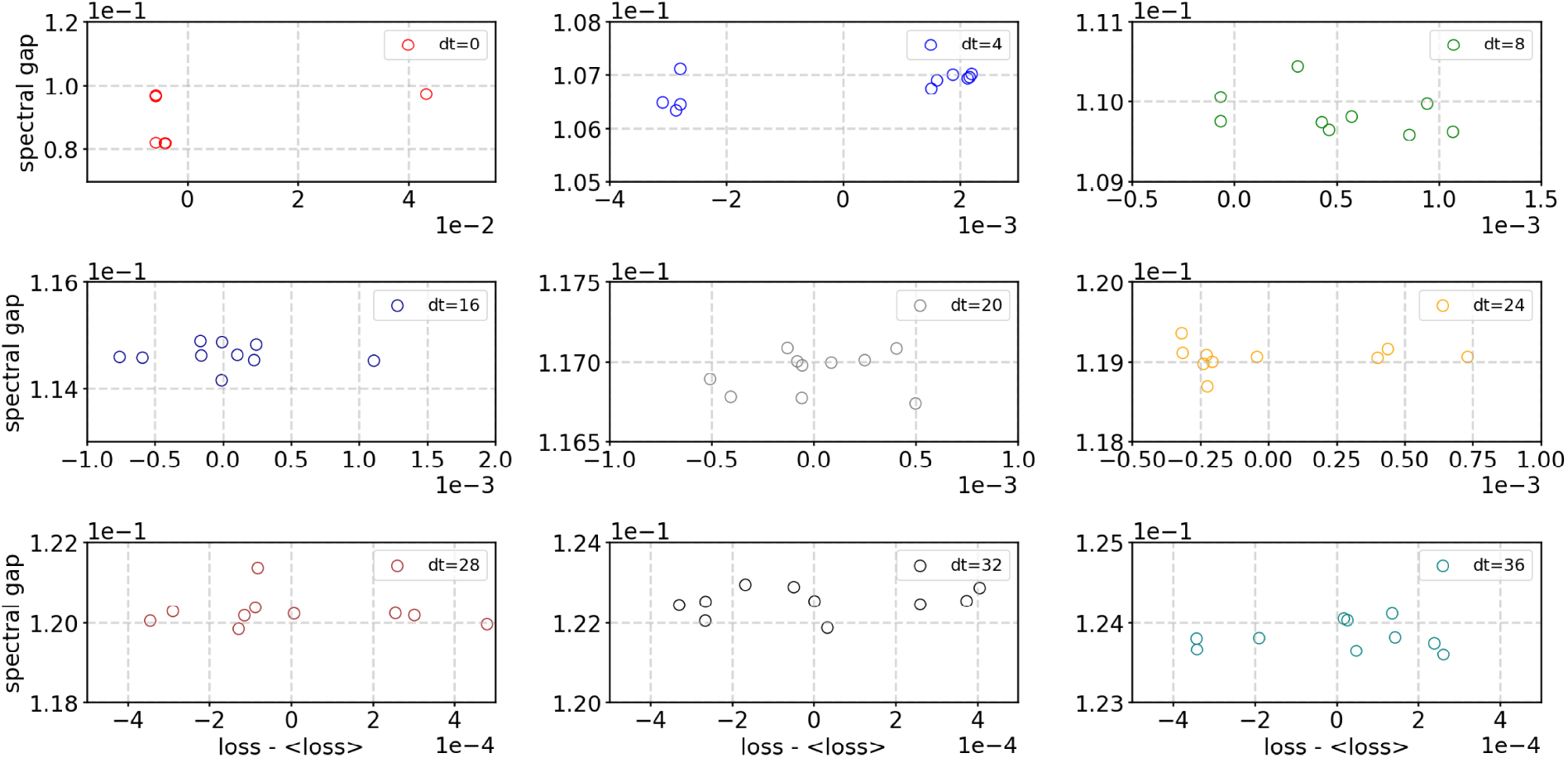
Loss function versus spectral gap for ligand unbinding from FKBP along the optimal reaction coordinate. Spectral gap and loss was calculated for the unbiased trajectory at multiple time-delays Δ*t* between 0 and 40 ps, indicated using circles of different colors. At each time-delay we observed a noisy correlation between the loss and the spectral gap. For visual clarity, we have plotted a mean-free version of the loss function by subtracting out the average of all losses.

**Figure S5:**
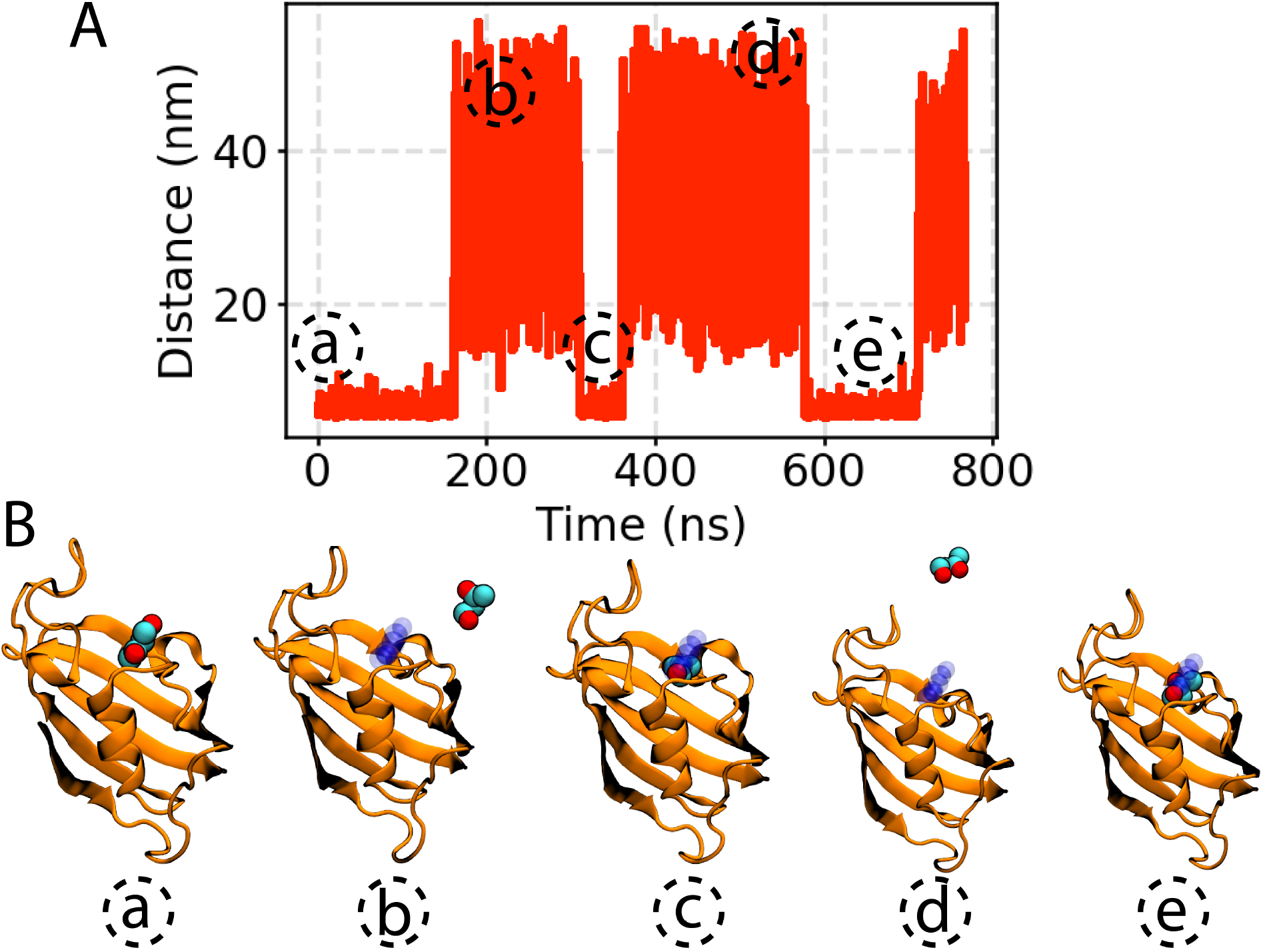
Biased simulation of ligand unbinding from FKBP along the optimal reaction coordinate. Biased simulation of ligand unbinding was performed under static bias. (A) Distance between the center of mass of the ligand from its binding pocket highlights multiple unbinding/binding events. (B) Representative snapshots of the trajectory are shown. Ligand is shown in vdW representation and the protein is shown in new cartoon (orange). Starting orientation of the ligand is shown in transparent blue vdW representation.

**Figure S6:**
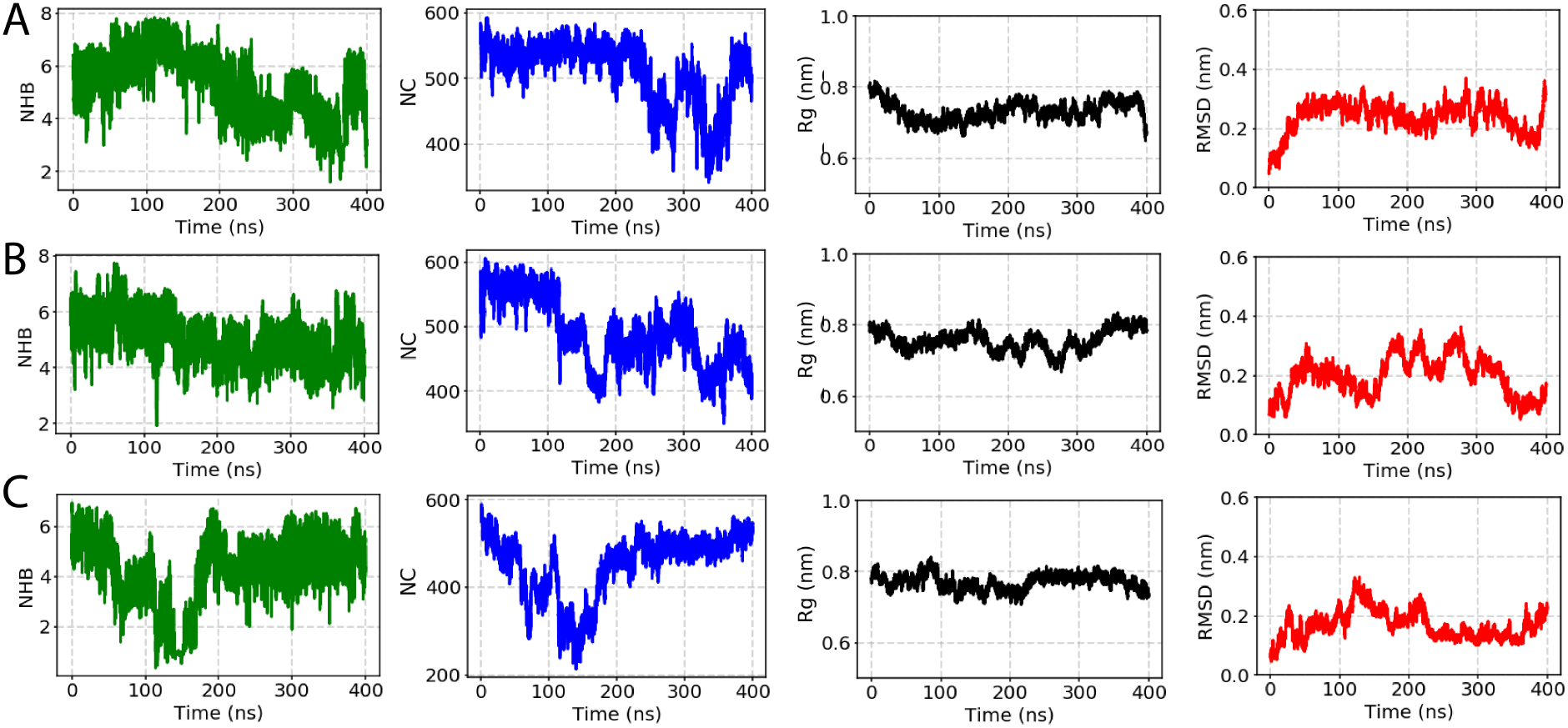
Unbiased MD simulations of GB1 peptide in explicit solvent. (A-C) Three independent MD simulations of GB1 peptide projected along different order parameters. Time evolution of OPs: root-mean square displacement (RMSD), radius of gyration (Rg), native contacts (NC), and number of backbone hydrogen bonds (NHB) suggests that the GB1 peptide was stable in all the simulated replicas.

**Figure S7:**
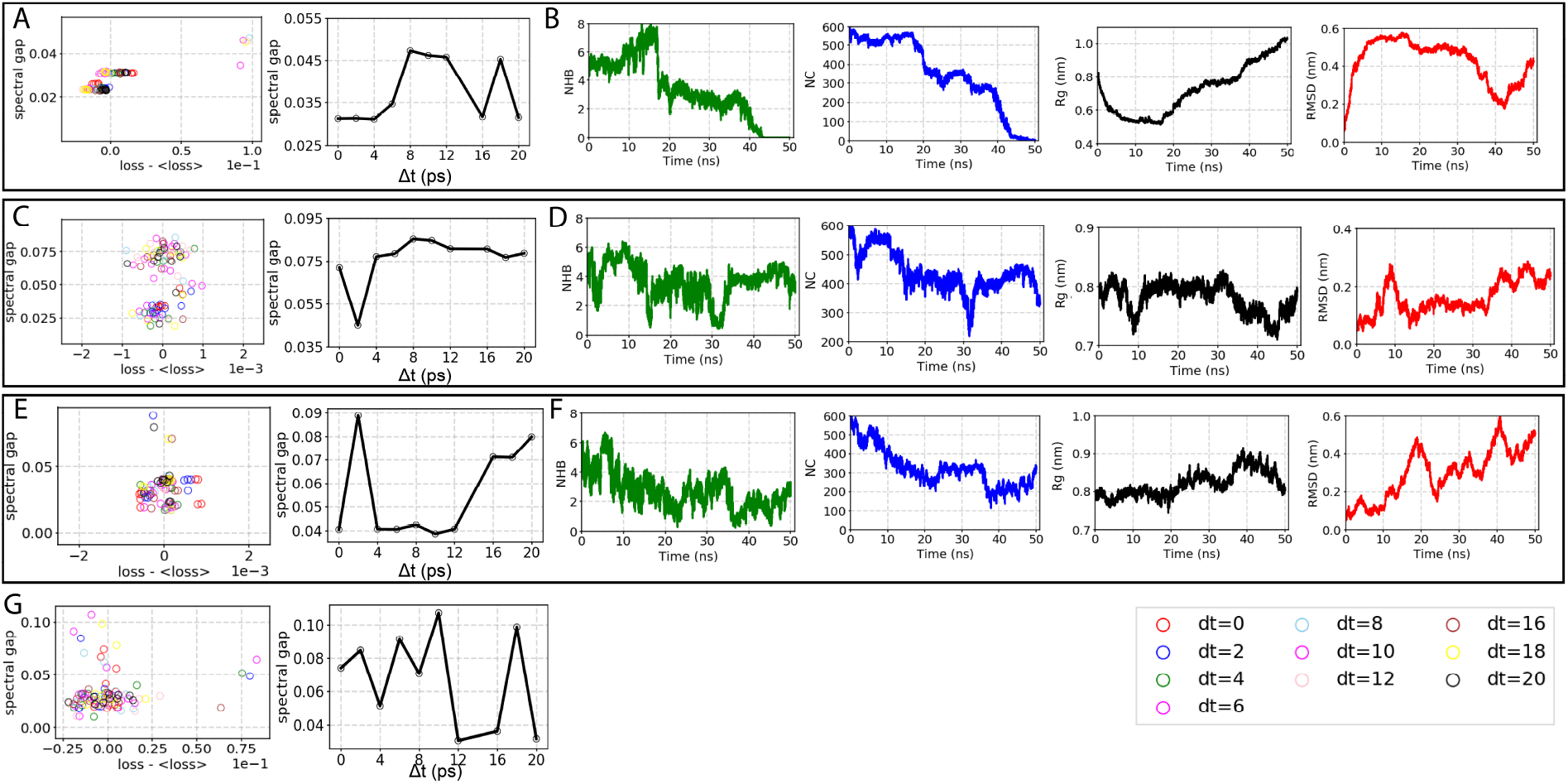
Capturing optimal reaction coordinate for GB1 peptide. Using an iterative ML-MD approach which incorporates RAVE and SGOOP, we have constructed an optimal RC to capture folding/unfolding of GB1 peptide. Spectral gap and loss was calculated in each round at multiple time-delays Δ*t* between 0 and 20 ps, indicated using circles of different colors. (A,C,E,G) Show noisy correlation between the loss and the spectral gap at each iteration. Different circles denote different independent trials, with color denoting Δ*t*. For visual clarity, at each iteration, we have plotted a mean-free version of the loss function by subtracting out the average of all losses. Corresponding maximum spectral gap (out of10 different trials of RAVE) vs. Δ*t* was plotted at each iteration. (B,D,F) 50 ns-long biased simulation along the constructed RC (at ieach iteration) was performed and the trajectory was projected along the order different parameters employed in this study.

**Figure S8:**
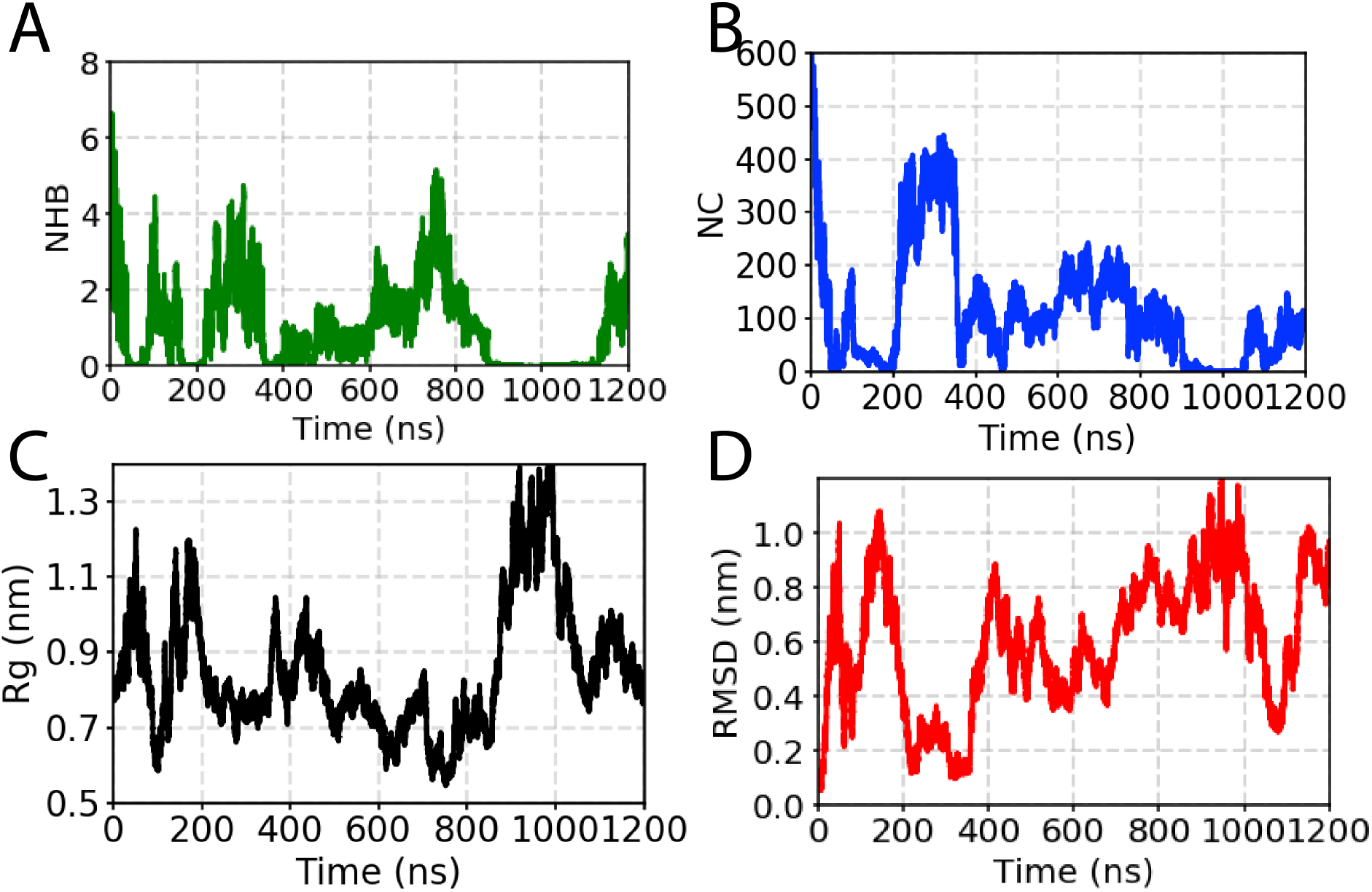
Biased simulation along the optimal 2-d reaction coordinate. The biased simulation was projected along the dictionary of order parameters used in this study. We captured multiple back and forth movement of the peptide between folded and unfolded states.

**Figure S9:**
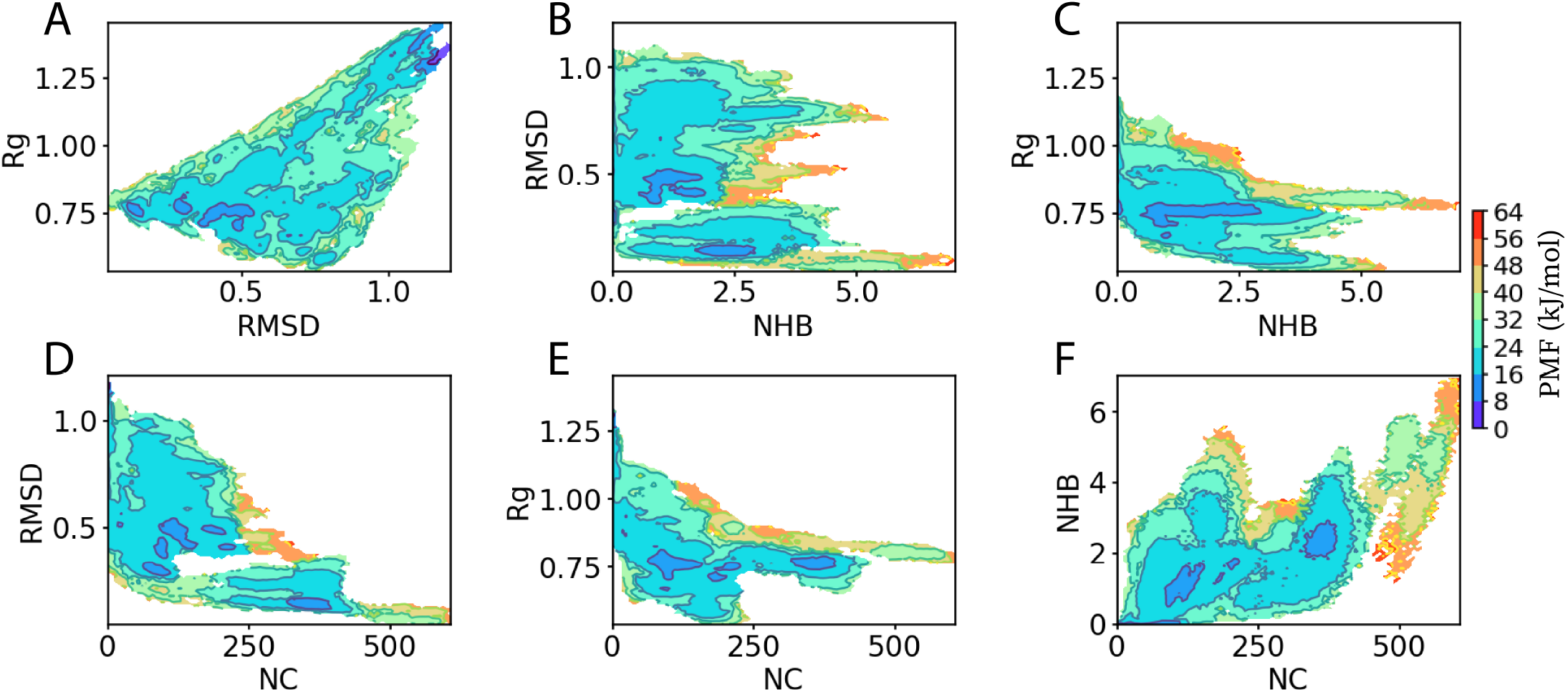
2-dimensional free-energy profiles along the different OPs: root-mean square displacement (RMSD), radius of gyration (Rg), native contacts (NC), and number of backbone hydrogen bonds (NHB) used in this study.

## Notes

### Competing Interest Statement

The authors have declared no competing interest.

## References

(1) Bernardi, R. C., Melo, M. C. R., Schulten, K. Enhanced Sampling Techniques in Molecular Dynamics Simulations of Biological Systems. Biochim. Biophys. Acta 2015, 1850, 872–877.

(2) Karplus, M., Petsko, G. A. Molecular dynamics simulations in biology. Nature 1990, 347, 631–639.

(3) Rohrdanz, M. A., Zheng, W., Clementi, C. Discovering mountain passes via torchlight: Methods for the definition of reaction coordinates and pathways in complex macromolecular reactions. Annu. Rev. Phys. Chem. 2013, 64, 295–316.

(4) Abrams, C., Bussi, G. Enhanced sampling in molecular dynamics using metadynamics, replica-exchange, and temperature-acceleration. Entropy 2014, 16, 163–199.

(5) Hashemian, B., Millán, D., Arroyo, M. Modeling and enhanced sampling of molecular systems with smooth and nonlinear data-driven collective variables. J. Chem. Phys. 2013, 139, 12B601 1.

(6) Onuchic, J. N., Wolynes, P. G. Theory of protein folding. Curr. Opin. Struct. Biol. 2004, 14, 70–75.

(7) Dill, K. A., Ozkan, S. B., Shell, M. S., Weikl, T. R. The protein folding problem. Annu. Rev. Biophys. 2008, 37, 289–316.

(8) Tiwary, P., Limongelli, V., Salvalaglio, M., Parrinello, M. Kinetics of protein–ligand unbinding: Predicting pathways, rates, and rate-limiting steps. Proc. Natl. Acad. Sci. USA 2015, 112, E386–E391.

(9) Tiwary, P., Mondal, J., Berne, B. J. How and when does an anticancer drug leave its binding site? Sci. Adv. 2017, 3, e1700014.

(10) Moradi, M., Tajkhorshid, E. Mechanistic picture for conformational transition of a membrane transporter at atomic resolution. Proc. Natl. Acad. Sci. USA 2013, 110, 18916–18921.

(11) Moradi, M., Tajkhorshid, E. Computational recipe for efficient description of large-scale conformational changes in biomolecular systems. J. Chem. Theory Comput. 2014, 10, 2866–2880.

(12) Moradi, M., Enkavi, G., Tajkhorshid, E. Atomic-level characterization of transport cycle thermodynamics in the glycerol-3-phosphate:phosphate transporter. Nat. Commun. 2015, 6, 8393.

(13) Pant, S., Tajkhorshid, E. Microscopic Characterization of GRP1 PH Domain Interaction with Anionic Membranes. J. Comput. Chem. 2019, 41, 489–499.

(14) Brooks, S. P., Morgan, B. J. Optimization using simulated annealing. J. of the Royal Statistical Society: Series D (The Statistician) 1995, 44, 241–257.

(15) Hansmann, U. H. Parallel tempering algorithm for conformational studies of biological molecules. Chemical Physics Letters 1997, 281, 140–150.

(16) Sugita, Y., Okamoto, Y. Replica-exchange molecular dynamics method for protein folding. Chem. Phys. Lett. 1999, 314, 141–151.

(17) Sugita, Y., Kitao, A., Okamoto, Y. Multidimensional replica-exchange method for free-energy calculations. J. Chem. Phys. 2000, 113, 6042–6051.

(18) Mitsutake, A., Sugita, Y., Okamoto, Y. Generalized-Ensemble Algorithms for Molecular Simulations of Biopolymers. Biopolymers 2001, 60, 96–123.

(19) Fukunishi, H., Watanabe, O., Takada, S. On the Hamiltonian replica exchange method for efficient sampling of biomolecular systems: Application to protein structure prediction. J. Chem. Phys. 2002, 116, 9058–9067.

(20) Maragliano, L., Vanden-Eijnden, E. A temperature accelerated method for sampling free energy and determining reaction pathways in rare events simulations. Chem. Phys. Lett. 2006, 426, 168–175.

(21) Miao, Y., Feher, V. A., McCammon, J. A. Gaussian accelerated molecular dynamics: Unconstrained enhanced sampling and free energy calculation. J. Chem. Theory Comput. 2015, 11, 3584–3595.

(22) Laio, A., Parrinello, M. Escaping free-energy minima. Proc. Natl. Acad. Sci. USA 2002, 99, 12562–12566.

(23) Laio, A., Gervasio, F. L.Metadynamics: a method to simulate rare events and reconstruct the free energy in biophysics, chemistry and material science. Rep. Progr. Phys. 2008, 71, 126601.

(24) Barducci, A., Bussi, G., Parrinello, M. Well-tempered metadynamics: a smoothly converging and tunable free-energy method. Phys. Rev. Lett. 2008, 100, 020603.

(25) Torrie, G. M., Valleau, J. P. Nonphysical sampling distributions in Monte Carlo free-energy estimation: Umbrella sampling. J. Chem. Phys. 1977, 23, 187–199.

(26) Lelièvre, T., Rousset, M., Stoltz, G. Computation of free energy profiles with parallel adaptive dynamics. J. Chem. Phys. 2007, 126, 134111.

(27) Darve, E., Rodríguez-Gómez, D., Pohorille, A. Adaptive biasing force method for scalar and vector free energy calculations. J. Chem. Phys. 2008, 128, 144120.

(28) Comer, J., Gumbart, J. C., Hénin, J., Lelièvre, T., Pohorille, A., Chipot, C. The adaptive biasing force method: Everything you always wanted to know but were afraid to ask. J. Phys. Chem. B 2015, 119, 1129– 1151.

(29) Fu, H., Shao, X., Chipot, C., Cai, W. Extended adaptive biasing force algorithm. An on-the-fly implementation for accurate free-energy calculations. J. Chem. Theory Comput. 2016, 12, 3506–3513.

(30) Lesage, A., Lelievre, T., Stoltz, G., Hénin, J. Smoothed biasing forces yield unbiased free energies with the extended-system adaptive biasing force method. J. Phys. Chem. B 2017, 121, 3676–3685.

(31) Abrams, J. B., Tuckerman, M. E. Efficient and direct generation of multidimensional free energy surfaces via adiabatic dynamics without coordinate transformations. J. Phys. Chem. B 2008, 112, 15742–15757.

(32) Kirkwood, J. G. Statistical mechanics of fluid mixtures. J. Chem. Phys. 1935, 3, 300–313.

(33) den Otter, W. K., Briels, W. J. The calculation of free-energy differences by constrained molecular-dynamics simulations. J. Chem. Phys. 1998, 109, 4139–4146.

(34) Hazel, A., Chipot, C., Gumbart, J. C. Thermodynamics of deca-alanine folding in water. J. Chem. Theory Comput. 2014, 10, 2836–2844.

(35) Liu, P., Kim, B., Friesner, R. A., Berne, B. Replica exchange with solute tempering: A method for sampling biological systems in explicit water. Proc. Natl. Acad. Sci. USA 2005, 102, 13749–13754.

(36) Huang, X., Hagen, M., Kim, B., Friesner, R. A., Zhou, R., Berne, B. J. Replica exchange with solute tempering: efficiency in large scale systems. J. Phys. Chem. B 2007, 111, 5405–5410.

(37) Wang, Y., Ribeiro, J. M. L., Tiwary, P. Machine learning approaches for analyzing and enhancing molecular dynamics simulations. Curr. Opin. Struct. Biol. 2020, 61, 139–145.

(38) Noé, F., De Fabritiis, G., Clementi, C. Machine learning for protein folding and dynamics. Curr. Opin. Struct. Biol. 2020, 60, 77–84.

(39) Noé, F., Olsson, S., Köhler, J., Wu, H. Boltzmann generators: Sampling equilibrium states of many-body systems with deep learning. Science 2019, 365, eaaw1147.

(40) Wang, Y., Ribeiro, J. M. L., Tiwary, P. Past–future information bottleneck for sampling molecular reaction coordinate simultaneously with thermodynamics and kinetics. Nat. Commun. 2019, 10, 1–8.

(41) Chen, W., Tan, A. R., Ferguson, A. L. Collective variable discovery and enhanced sampling using autoencoders: Innovations in network architecture and error function design. J. Comput. Chem. 2018, 149, 072312.

(42) Tuckerman, M. E. Machine learning transforms how microstates are sampled. Science 2019, 365, 982–983.

(43) Lahey, S.-L. J., Rowley, C. N. Simulating protein–ligand binding with neural network potentials. Chem. Sci. 2020, 11, 2362–2368.

(44) Rotskoff, G., Vanden-Eijnden, E. Parameters as interacting particles: long time convergence and asymptotic error scaling of neural networks. Advances in neural information processing systems. 2018; pp 7146–7155.

(45) Cybenko, G. Approximation by superpositions of a sigmoidal function. Mathematics of Control, Signals and Systems 1992, 5, 455–455.

(46) Barron, A. R. Universal approximation bounds for superpositions of a sigmoidal function. IEEE Transactions on Information theory 1993, 39, 930–945.

(47) Bach, F. Breaking the curse of dimensionality with convex neural networks. The Journal of Machine Learning Research 2017, 18, 629–681.

(48) Evtimov, I., Eykholt, K., Fernandes, E., Kohno, T., Li, B., Prakash, A., Rahmati, A., Song, D. Robust physical-world attacks on deep learning models. arXiv preprint arXiv:1707.08945 2017,

(49) Noé, F., Horenko, I., Schütte, C., Smith, J. C. Hierarchical analysis of conformational dynamics in biomolecules: Transition networks of metastable states. J. Chem. Phys. 2007, 126, 04B617.

(50) Rohrdanz, M. A., Zheng, W., Maggioni, M., Clementi, C. Determination of reaction coordinates via locally scaled diffusion map. J. Chem. Phys. 2011, 134, 03B624.

(51) Noé, F., Nuske, F. A variational approach to modeling slow processes in stochastic dynamical systems. Multiscale Modeling & Simulation 2013, 11, 635–655.

(52) Pérez-Hernández, G., Paul, F., Giorgino, T., De Fabritiis, G., Noé, F. Identification of slow molecular order parameters for Markov model construction. J. Chem. Phys. 2013, 139, 07B604 1.

(53) Li, Q., Dietrich, F., Bollt, E. M., Kevrekidis, I. G. Extended dynamic mode decomposition with dictionary learning: A data-driven adaptive spectral decomposition of the Koopman operator. Chaos: An Interdisciplinary Journal of Nonlinear Science 2017, 27, 103111.

(54) Tiwary, P., Berne, B. Spectral gap optimization of order parameters for sampling complex molecular systems. Proc. Natl. Acad. Sci. USA 2016, 113, 2839–2844.

(55) Ghosh, K., Dixit, P. D., Agozzino, L., Dill, K. A. The Maximum Caliber Variational Principle for Nonequilibria. Annu. Rev. Phys. Chem. 2020, 71, 213–238.

(56) Ribeiro, J. M. L., Bravo, P., Wang, Y., Tiwary, P. Reweighted autoencoded variational Bayes for enhanced sampling (RAVE). J. Chem. Phys. 2018, 149, 072301.

(57) Ravindra, P., Smith, Z., Tiwary, P. Automatic mutual information noise omission (AMINO): generating order parameters for molecular systems. Molecular Systems Design & Engineering 2020,

(58) Tishby, N., Pereira, F. C., Bialek, W. The information bottleneck method. arXiv preprint physics/0004057 2000,

(59) Palmer, S. E., Marre, O., Berry, M. J., Bialek, W. Predictive information in a sensory population. Proc. Natl. Acad. Sci. USA 2015, 112, 6908–6913.

(60) Berman, G. J., Bialek, W., Shaevitz, J. W. Predictability and hierarchy in Drosophila behavior. Proc. Natl. Acad. Sci. USA 2016, 113, 11943–11948.

(61) Alemi, A. A., Fischer, I., Dillon, J. V., Murphy, K. Deep variational information bottleneck. arXiv preprint arXiv:1612.00410 2016,

(62) Still, S. Information bottleneck approach to predictive inference. Entropy 2014, 16, 968–989.

(63) Krivov, S. V. On reaction coordinate optimality. Journal of chemical theory and computation 2013, 9, 135– 146.

(64) Smith, Z., Ravindra, P., Wang, Y., Cooley, R., Tiwary, P. Discovering loop conformational flexibility in T4 lysozyme mutants through Artificial Intelligence aided Molecular Dynamics. bioRxiv 2020,

(65) Valsson, O., Tiwary, P., Parrinello, M. Enhancing important fluctuations: Rare events and metadynamics from a conceptual viewpoint. 2016, 67, 159–184.

(66) Cover, T. M., Thomas, J. A. Elements of information theory; John Wiley & Sons, 2012.

(67) Goodfellow, I., Bengio, Y., Courville, A. Deep Learning; MIT Press, 2016; http://www.deeplearningbook.org.

(68) Wang, Y., Tiwary, P. Understanding the role of predictive time delay and biased propagator in RAVE. J. Chem. Phys. 2020, 152, 144102.

(69) Levine, I. N., Busch, D. H., Shull, H. Quantum chemistry; Pearson Prentice Hall Upper Saddle River, NJ, 2009; Vol. 6.

(70) Nelson, D. L., Lehninger, A. L., Cox, M. M. Lehninger principles of biochemistry; Macmillan, 2008.

(71) Truhlar, D. G., Garrett, B. C. Variational transition state theory. Annu. Rev. Phys. Chem. 1984, 35, 159–189.

(72) Dixit, P. D., Dill, K. A. Caliber corrected Markov modeling (C2M2): Correcting equilibrium Markov models. J. Chem. Theory Comput. 2018, 14, 1111–1119.

(73) Smith, Z., Pramanik, D., Tsai, S.-T., Tiwary, P. Multi-dimensional spectral gap optimization of order parameters (SGOOP) through conditional probability factorization. J. Chem. Phys. 2018, 149, 234105.

(74) Meral, D., Provasi, D., Filizola, M. An efficient strategy to estimate thermodynamics and kinetics of G protein-coupled receptor activation using metadynamics and maximum caliber. J. Chem. Phys. 2018, 149, 224101.

(75) Tiwary, P., van de Walle, A. Multiscale Materials Modeling for Nanomechanics; Springer, 2016; pp 195– 221.

(76) Debnath, J., Parrinello, M. Gaussian Mixture Based Enhanced Sampling For Statics And Dynamics. J. Phys. Chem. Lett. 2020,

(77) Abraham, M. J., Murtola, T., Schulz, R., Páll, S., Smith, J. C., Hess, B., Lindahl, E. GROMACS: High performance molecular simulations through multi-level parallelism from laptops to supercomputers. SoftwareX 2015, 1, 19–25.

(78) Tribello, G. A., Bonomi, M., Branduardi, D., Camilloni, C., Bussi, G. PLUMED 2: New feathers for an old bird. Comput. Phys. Commun. 2014, 185, 604–613.

(79) Bonomi, M., Bussi, G., Camilloni, C., Tribello, G. A., Banáš, P., Barducci, A., Bernetti, M., Bolhuis, P. G., Bottaro, S., Branduardi, D., et al. Promoting transparency and reproducibility in enhanced molecular simulations. Nat. Methods

(80) Mu, Y., Nguyen, P. H., Stock, G. Energy landscape of a small peptide revealed by dihedral angle principal component analysis. Proteins: Struct., Func., Bioinf. 2005, 58, 45–52.

(81) Altis, A., Nguyen, P. H., Hegger, R., Stock, G. Dihedral angle principal component analysis of molecular dynamics simulations. J. Chem. Phys. 2007, 126, 244111.

(82) Salvalaglio, M., Tiwary, P., Parrinello, M. Assessing the reliability of the dynamics reconstructed from metadynamics. J. Chem. Theory Comput. 2014, 10, 1420–1425.

(83) Hornak, V., Abel, R., Okur, A., Strockbine, B., Roitberg, A., Simmerling, C. Comparison of multiple Amber force fields and development of improved protein backbone parameters. Proteins: Struct., Func., Bioinf. 2006, 65, 712–725.

(84) Best, R. B., Hummer, G. Optimized molecular dynamics force fields applied to the helix-coil transition of polypeptides. J. Phys. Chem. B 2009, 113, 9004–9015.

(85) Lindorff-Larsen, K., Piana, S., Palmo, K., Maragakis, P., Klepeis, J. L., Dror, R. O., Shaw, D. E. Improved side-chain torsion potentials for the Amber ff99SB protein force field. Proteins: Struct., Func., Bioinf. 2010, 78, 1950–1958.

(86) Gumbart, J. C., Roux, B., Chipot, C. Standard binding free energies from computer simulations: What is the best strategy? Journal of chemical theory and computation 2013, 9, 794–802.

(87) Burkhard, P., Taylor, P., Walkinshaw, M. D. X-ray structures of small ligand-FKBP complexes provide an estimate for hydrophobic interaction energies. J. Mol. Biol. 2000, 295, 953–962.

(88) Pramanik, D., Smith, Z., Kells, A., Tiwary, P. Can One Trust Kinetic and Thermodynamic Observables from Biased Metadynamics Simulations?: Detailed Quantitative Benchmarks on Millimolar Drug Fragment Dissociation. J. Phys. Chem. B 2019, 123, 3672–3678.

(89) Pan, A. C., Xu, H., Palpant, T., Shaw, D. E. Quantitative characterization of the binding and unbinding of millimolar drug fragments with molecular dynamics simulations. Journal of chemical theory and computation 2017, 13, 3372–3377.

(90) Ahalawat, N., Mondal, J. Assessment and optimization of collective variables for protein conformational landscape: GB1 β-hairpin as a case study. J. Chem. Phys. 2018, 149, 094101.

(91) Munoz, V., Thompson, P. A., Hofrichter, J., Eaton, W. A. Folding dynamics and mechanism of β-hairpin formation. Nature 1997, 390, 196–199.

(92) Fesinmeyer, R. M., Hudson, F. M., Andersen, N. H. Enhanced hairpin stability through loop design: the case of the protein G B1 domain hairpin. J. Am. Chem. Soc. 2004, 126, 7238–7243.

(93) Hazel, A. J., Walters, E. T., Rowley, C. N., Gumbart, J. C. Folding free energy landscapes of β-sheets with non-polarizable and polarizable CHARMM force fields. J. Chem. Phys. 2018, 149, 072317.

(94) Best, R. B., Mittal, J. Free-energy landscape of the GB1 hairpin in all-atom explicit solvent simulations with different force fields: Similarities and differences. Proteins: Struct., Func., Bioinf. 2011, 79, 1318–1328.

(95) Ardevol, A., Tribello, G. A., Ceriotti, M., Parrinello, M. Probing the unfolded configurations of a β-hairpin using sketch-map. J. Chem. Theory Comput. 2015, 11, 1086–1093.

(96) Nuske, F., Keller, B. G., Pérez-Hernández, G., Mey, A. S., Noé, F. Variational approach to molecular kinetics. J. Chem. Theory Comput. 2014, 10, 1739–1752.

(97) McGibbon, R. T., Pande, V. S. Variational cross-validation of slow dynamical modes in molecular kinetics. J. Chem. Phys. 2015, 142, 03B621 1.

(98) Lamim Ribeiro, J. M., Tiwary, P. Toward achieving efficient and accurate Ligand-Protein unbinding with deep learning and molecular dynamics through RAVE. J. Chem. Theory Comput. 2018, 15, 708–719.

(99) Bonomi, M., Branduardi, D., Gervasio, F. L., Parrinello, M. The unfolded ensemble and folding mechanism of the C-terminal GB1 β-hairpin. J. Am. Chem. Soc. 2008, 130, 13938–13944.

(100) Bussi, G., Gervasio, F. L., Laio, A., Parrinello, M. Free-energy landscape for β hairpin folding from combined parallel tempering and metadynamics. J. Am. Chem. Soc. 2006, 128, 13435–13441.

(101) Saladino, G., Pieraccini, S., Rendine, S., Recca, T., Francescato, P., Speranza, G., Sironi, M. Metadynamics study of a β-hairpin stability in mixed solvents. J. Am. Chem. Soc. 2011, 133, 2897–2903.

(102) Pressé, S., Ghosh, K., Lee, J., Dill, K. A. Principles of maximum entropy and maximum caliber in statistical physics. Rev. Mod. Phys. 2013, 85, 1115.

(103) Chen, W., Sidky, H., Ferguson, A. L. Nonlinear discovery of slow molecular modes using state-free reversible VAMPnets. J. Chem. Phys. 2019, 150, 214114.

(104) Bussi, G., Donadio, D., Parrinello, M. Canonical sampling through velocity rescaling. J. Chem. Phys. 2007, 126, 014101.

(105) Wang, J., Wolf, R. M., Caldwell, J. W., Kollman, P. A., Case, D. A. Development and testing of a general amber force field. J. Comput. Chem. 2004, 25, 1157–1174.

(106) Martyna, G. J., Tobias, D. J., Klein, M. L. Constant pressure molecular dynamics algorithms. J. Chem. Phys. 1994, 101, 4177–4189.

(107) Tiwary, P., Parrinello, M. A time-independent free energy estimator for metadynamics. J. Phys. Chem. B 2015, 119, 736–742.

